# Extrachromosomal Telomeres Derived from Excessive Strand Displacements

**DOI:** 10.1101/2023.07.31.551186

**Authors:** Junyeop Lee, Jina Lee, Eric J. Sohn, Angelo Taglialatela, Roderick J. O’Sullivan, Alberto Ciccia, Jaewon Min

## Abstract

Alternative Lengthening of Telomeres (ALT) is a telomere maintenance mechanism mediated by break-induced replication (BIR), evident in approximately 15% of human cancers. A characteristic feature of ALT cancers is the presence of C-circles, circular single-stranded telomeric DNAs composed of C-rich sequences. Despite the fact that extrachromosomal C-rich single-stranded DNAs (ssDNAs), unique to ALT cells, are considered potential precursors of C-circles, their generation process remains undefined. Here, we introduce a highly sensitive method to detect single stranded telomeric DNA, called 4SET (Strand-Specific Southern-blot for Single-stranded Extrachromosomal Telomeres) assay. Utilizing 4SET, we are able to capture C-rich single stranded DNAs that are near 200 to 1500 nucleotides in size. Both linear C-rich ssDNAs and C-circles are abundant in the fractions of cytoplasm and nucleoplasm, which supports the idea that linear C-rich ssDNA accumulation may indeed precede C-circle formation. We also found that C-rich ssDNAs originate during Okazaki fragment processing during lagging strand DNA synthesis. The generation of C-rich ssDNA requires CST-PP (CTC1/STN1/TEN1-PRIMASE-Polymerase alpha) complex-mediated priming of the C-strand DNA synthesis and subsequent excessive strand displacement of the C-rich strand mediated by the DNA Polymerase delta and the BLM helicase. Our work proposes a new model for the generation of C-rich ssDNAs and C-circles during ALT-mediated telomere elongation.

## Introduction

Telomeres are repetitive sequences consisting of TTAGGG at the ends of chromosomes (1). With each cell division, telomeres undergo progressive shortening due to the end replication problem. Consequently, during tumorigenesis, cancer cells acquire a mechanism to maintain telomere length in order to undergo neoplastic transformation. While most cancers activate telomerase, a reverse transcriptase which synthesizes telomere sequences, a subset of cancers acquire a mechanism called alternative lengthening of telomeres (ALT), which operates independently of telomerase and elongates telomeres through a DNA recombination-dependent process known as break-induced replication (BIR) (2–6). ALT cancers are characterized by ALT-associated biomarkers, including ALT-associated PML bodies (APBs) and extrachromosomal telomeric circular DNAs, such as C-circles (Cytosine-rich sequences (CCCTAA) containing single stranded circular telomeric DNAs) and T-circles (t-loop sized circle; double stranded circular telomeric DNAs) (7).

T-circles, which can be visualized using electron microscopy or high-resolution STORM microscopy, are generated through processes involving telomere trimming, I-loop (internal-loop) resolution, and looping out during replication processes (8–12). One well-characterized mechanism for T-circle generation is the involvement of TZAP, a negative regulator of telomere length implicated in telomere trimming (13). Notably, T-circles are not only abundant in ALT cancer cells but also present in telomerase-positive cells and germ cells, which possess very long telomeres (14, 15). Notably, these T-circles are generated in PML bodies (12).

C-circles were initially found in ALT cancer and ALT-positive immortalized cell lines (16), making them valuable diagnostic markers for ALT cancer (17–19). These circular DNA molecules are present in low abundance and are mostly detectable through the use of phi29 polymerase (16). Interestingly, recent observations have shown that C-circles are also present in very long telomere-containing fibroblasts, even in cases where ALT is not involved (20). Furthermore, C-circles seem to preferentially form from the lagging strand (21, 22). The generation of C-circles in ALT cancer cells is thought to be related to replication stress experienced during S phases (23). Replication stress potentially activates the ALT pathway and leads to the formation of C-circles. However, the precise process by which C-circles are generated in ALT cancer cells remains unclear, necessitating further research to uncover the underlying mechanisms.

Apart from telomeric circular DNAs, ALT telomeres display various other structures, including T-bands, 5’ overhangs, and single-stranded telomeric DNAs (24, 25). T-bands and 5’ overhangs are predominantly observed in the chromatin fraction (25). The presence of 5’ overhangs is likely a consequence of telomere trimming, as they can be detected during T-circle generation processes (15). Notably, a recent study has revealed that T-bands are derived from template switching during BIR-mediated ALT telomere synthesis (26).

Single-stranded telomeric DNAs, on the other hand, are extrachromosomal structures that are enriched in the cytoplasmic fraction obtained by Hirt Lysate (24, 25). These single-stranded extrachromosomal telomeric DNAs are believed to serve as potential precursors for C-circles. However, the specific process through which extrachromosomal C-rich single-stranded telomere DNA generation occurs remains unclear and requires further investigation to understand its underlying mechanisms.

In this study, we developed the 4SET (Strand-Specific Southern-blot for Single-stranded Extrachromosomal Telomeres) assay, a simple and efficient method for detecting these telomeres. Using 4SET, we are able to capture telomeric DNA fragments that contain C-rich single stranded strands of DNA that can be easily measured, around 200 to 1500 nucleotides in size. Both C-rich ssDNAs and C-circles are abundant in the cytoplasmic fraction, which supports the idea that linear C-rich ssDNAs are potential precursors of C-circles. We also found that C-rich ssDNAs are generated during Okazaki fragment processing in lagging strand DNA synthesis. The generation of C-rich ssDNA requires CST-PP (CTC1/STN1/TEN1-PRIMASE-Polymerase alpha) complex-mediated priming of the C-strand DNA synthesis and subsequent excessive strand displacement of the C-rich strand mediated by the DNA Polymerase delta. Our work proposes a new model for the generation of C-rich ssDNAs and C-circles, which are markers of an active ALT pathway.

## Results

### 4SET assay, a simple and efficient method for detecting single-stranded extrachromosomal telomeric DNA

To investigate the abundance of single-stranded telomeric DNA in ALT cancer, we have made modifications to existing Southern protocols that we named 4SET (Strand-Specific Southern-blot for Single-stranded Extrachromosomal Telomeres) (Fig. 1A). These modifications enhance sensitivity and optimize detection of single-stranded DNA. First, we optimized the genomic DNA extraction procedure for single-stranded DNA fragments. It has been observed that the isopropanol/ethanol precipitation procedure can result in unreliable nucleotides recovery efficiency, particularly for small nucleic acids when the concentration of nucleic acid is low (27). During isopropanol/ethanol DNA precipitation, the presence of bulk DNA/RNA can act as a carrier, which facilitates the precipitation of small-sized nucleic acids. To overcome this issue, we introduced excessive amounts of glycogen as a carrier during the isopropanol precipitation step. Additionally, we incubated the mixture at −20 degrees Celsius for a minimum of 3 hours to maximize the precipitation of single-stranded DNA fragments. Compared to the conventional genomic DNA extraction kit (Qiagen DNeasy), our method is specifically optimized for the preparation of small-sized nucleic acids (Fig. S1A). Secondly, we ran the genomic DNA on a low-percentage agarose gel and conducted the transfer onto a positively charged nylon membrane instead of using gel dry. This approach was chosen primarily to avoid the loss of small DNA fragments that can occur during the vacuum-based gel drying process. When the gel is dried using a vacuum, there is a risk of small DNA fragments being lost as the moisture is removed from below to dry the gel through the pressure. Lastly, we used highly sensitive strand specific C-rich and G-rich telomere probe to determine strand specificity (28). This probe is particularly optimal for detecting C-rich ssDNAs compared to probes based on end-labeling (Fig. S1B).

**Figure 1.**
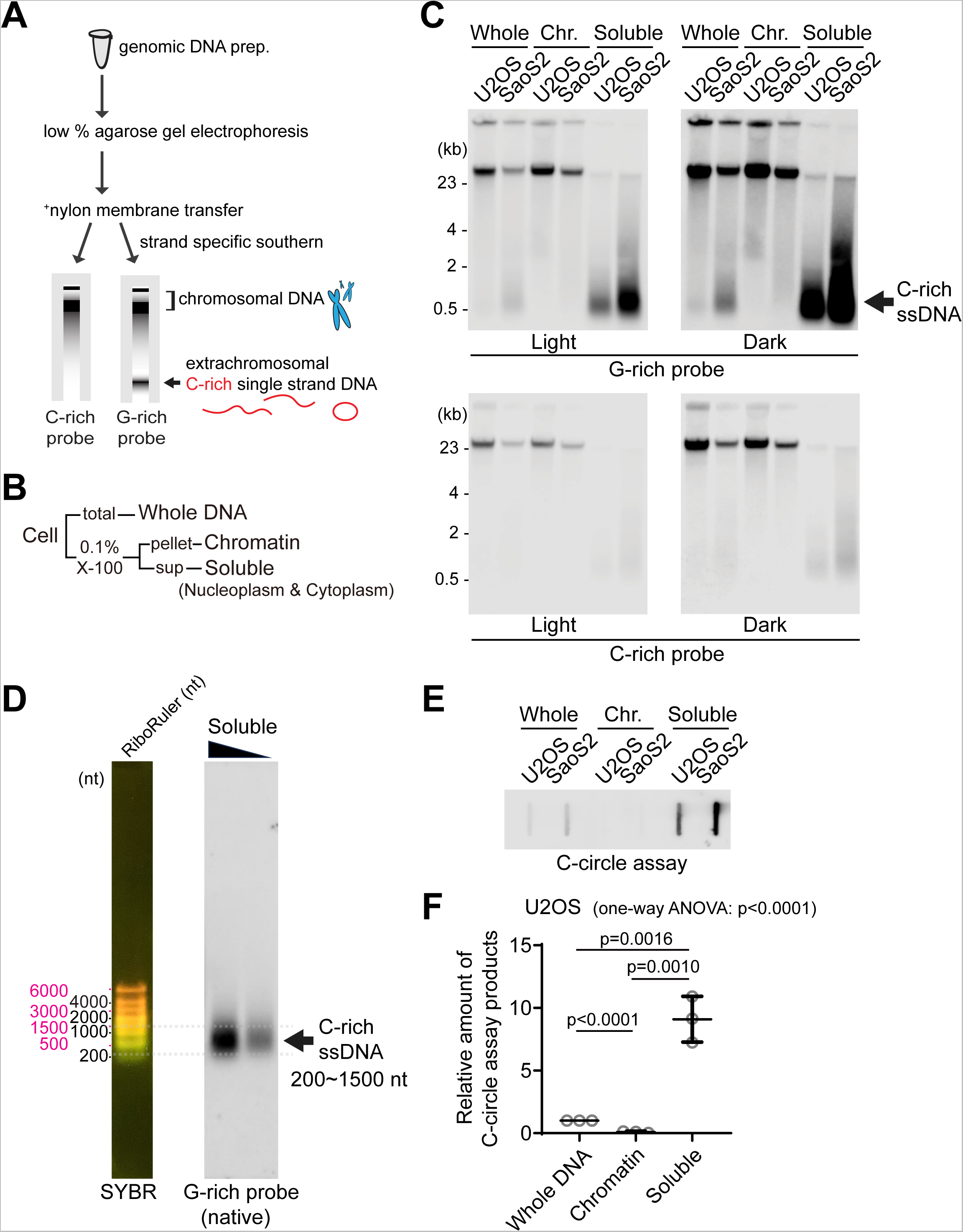
Detection of C-rich extrachromosomal telomeres. **(A)** Illustration of the Strand-specific Southern for Single Stranded Extrachromosomal Telomeres (4SET) method. (Refer to the method for the details) **(B)** The 4SET method involves the fractionation of samples into Whole DNA fraction, Chromatin fraction, and Triton X-100 soluble fraction, achieved using 0.1% Triton X-100 extraction buffer. **(C)** 4SET assay for U2OS and SaoS2 cells after fractionation of samples into Whole, Chromatin (Chr.), and Soluble fractions. DNA amount: 600 ng. G-rich probe was used to detect C-rich telomeric sequences, and C-rich probe was used to detect G-rich telomeric sequences. C-rich single stranded DNAs are indicated by an arrow. **(D)** 4SET assay with a nucleotide ladder (RiboRuler) to determine the size of C-rich single-stranded DNA in SaoS2 soluble fraction. G-rich probe was used for native gel to detect C-rich single stranded telomeres. **(E)** C-circle assay for U2OS and SaoS2 cells after fractionation of samples into Whole, Chromatin, and Soluble fractions. DNA amount: 50ng **(F)** Quantification of the C-circle assay in E; U2OS cells as the relative amount of C-circle assay products (mean ± SD; unpaired t-test), One-way ANOVA analysis; p<0.0001.

Furthermore, we carefully considered the temperature at which we dissolve the precipitated DNA pellet. Long incubation at high temperatures, such as 55 degrees Celsius, is not optimal for the assay (Fig. S1C). As previously reported, C-circles are also sensitive to temperature and freeze-thaw cycle (16). DNA dissolution temperature also affects C-circle assay due to its stability. We thus used low temperatures for DNA dissolution and avoided freeze-thaw cycle.

Single-stranded telomeres are separated via Hirt lysate, a method used for isolating extrachromosomal DNA (24, 25). Consequently, we conducted cell fractionation to investigate the presence of C-rich ssDNAs in the cytoplasm. By employing 0.1% Triton X-100, we separated the cells into a chromatin fraction (pellet) and a soluble fraction (supernatant; nucleoplasm and cytoplasm) (Fig. 1B). We validated our cell fractionation by Western blot, confirming that histone H3 proteins are present in chromatin and whole fractions, GAPDH proteins are found in soluble and whole fractions, and beta-actin proteins are enriched in soluble fractions (Fig. S1D).

We performed 4SET assay after cell fractionation in U2OS and SaoS2 cells, which are commonly used ALT-positive cancer cell lines derived from osteosarcoma patients (Fig. 1C, S1E). As previous reports showed (24), SaoS2 cells exhibited bands around 500 bp in size that correspond to C-rich ssDNA (Fig. 1C; G-rich probe). Notably, the cytoplasmic fraction exhibited a substantial abundance of C-rich ssDNAs, while no signal was detected in the chromatin fraction, indicating that C-rich ssDNA predominantly resides in the supernatant (Fig. 1D; soluble DNA). Furthermore, our 4SET method enabled us to detect these small single stranded C-rich telomeres in U2OS cells as well, which were not detected previously (Fig. 1C; U2OS soluble DNA).

Extrachromosomal telomeric circular DNAs are thought to be generated in ALT-associated PML bodies (APBs). We detected that PML bodies existed in both chromatin and soluble fractions (Fig. S1F). To determine if PML bodies are responsible for the generation of C-rich ssDNA, we generated PML knockout in SaoS2 cells using CRISPR/Cas9 (Fig. S1G). PML knockout SaoS2 cells do not harbor C-rich ssDNAs (Fig. S1H), suggesting that C-rich ssDNAs are generated in PML bodies. Next, we measured the C-circles by using phi29 polymerase mediated rolling circle amplification; PML knockout cells also harbor less abundant level of C-circles (Fig. S1I).

We determined the size of the C-rich ssDNA using the nucleotide ladder (RiboRuler). The sizes of the C-rich ssDNAs ranged from approximately 200 to 1500 nt (Fig. 1D). Their size distribution aligns more closely with the RiboRuler than with the base pair ladder (GeneRuler) when run on both 0.5% and 1% native agarose gels (Fig. S1J), supporting that these are single-stranded DNAs.

Induction of DNA double-strand breaks using Zeocin, Etoposide (ETO), and Camptothecin (CPT) produced signals exceeding 23 kb in the soluble fraction (Fig. S1K). This likely represents broken chromosomes released into the soluble fraction, supporting the idea that the soluble fraction contains extrachromosomal DNAs. Notably, while treatment with the Topoisomerase 1 inhibitor CPT led to an increase in C-rich ssDNA generation, Zeocin treatment, which induces DNA double-strand breaks, did not have this effect. This suggests that C-rich ssDNAs are produced during DNA replication or transcription, rather than being a direct result of inducing DNA cleavages or DNA double-strand breaks at telomeres.

Next, we measured the C-circles after cell fractionation (Fig. 1E). C-circles were also detected more in the soluble fraction (Fig. 1F, S1L), which is in line with original findings on C-circles (16). We employed an alternative conventional method to fractionate cells using E1 and E2 buffers. This approach separates the cells into chromatin, nucleoplasm, and cytoplasm fractions (Fig. S1M-N). We observed that C-rich ssDNAs are enriched in both the nucleoplasm and cytoplasm fractions (Fig. S1O) which aligns with the soluble fraction enrichment data (Fig. 1C).

### Originating from lagging daughter strands, C-rich ssDNAs contain both linear and circular DNAs

We conducted an S1 endonuclease digestion which specifically cleaves ssDNA. All signals in soluble fraction disappeared upon S1 nuclease treatment confirming that the those are indeed ssDNA (Fig. S2A). To determine whether these are linear or circular DNAs, we performed exonuclease digestion (Fig. 2A). Approximately 50% of the signals were resistant to exonucleases, suggesting that they could be circular DNAs (Fig. 2B, S2B). In addition to the 4SET assay, C-rich ssDNAs in soluble fractions were confirmed as single-stranded DNA using a telomere-PNA-probe through native telomere-FISH (Fig. S2C).

**Figure 2.**
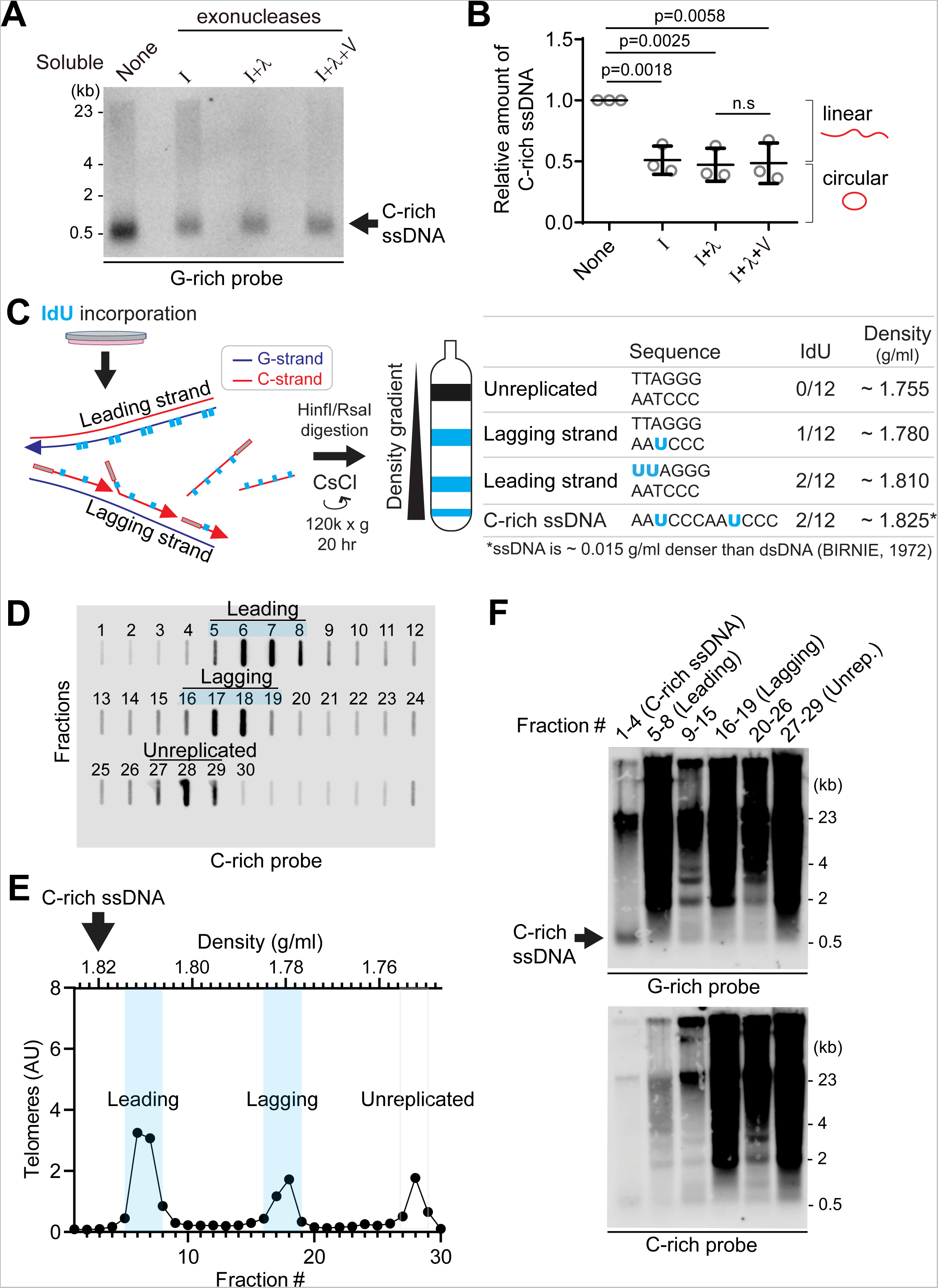
C-rich ssDNAs from lagging daughter strands contain both linear and circular DNA. **(A)** Nuclease assay using exonuclease I, lambda, and V treatment on soluble DNA fraction of U2OS. 180 ng of soluble fraction DNA was used per lane. **(B)** Quantification of the nuclease assay in A; Relative amount of C-rich ssDNA (mean ± SD; unpaired t-test). **(C)** Illustration of the Iododeoxyuridine (IdU) incorporation and CsCl separation. The right table displays the expected density of unreplicated DNA and IdU-incorporated lagging strands, leading strands, and nascent C-rich single-stranded DNAs. **(D)** Slot blot analysis of telomeres in CsCl separation fractions. 150 µg of genomic DNA from U2OS cells was used. **(E)** Density and relative telomere intensities of fractions in D. **(F)** 4SET assay of the fractions depicted in D and E.

To elucidate the origin of C-rich ssDNAs, we utilized Cesium density gradient analysis (CsCl separation), a method originally used for differentiating leading and lagging strand telomeres. The incorporation of Iodo-deoxyuridine (IdU) leads to distinct density differences between these strands due to IdU’s denser nature compared to thymidine (Fig. 2C). We hypothesized that if C-rich ssDNAs originate from lagging daughter strands, their expected density (around 1.825 g/ml) would be slightly denser than the leading strand (approximated at 1.810 g/ml) (Fig. 2C). This is because ssDNA is about 0.015 g/ml denser than dsDNA (29), despite the fact that IdU-incorporated C-rich ssDNAs and leading telomeres have same IdU-incorporated composition (2 IdU per 12 nucleotide) (Fig. 2C; right table). After performing CsCl separation on the genomic DNA of U2OS cells, we were able to distinguish between the leading, lagging, and unreplicated DNA fractions based on their density (Fig. 2D-E), as we showed in our previous reports (21). Confirming our hypothesis, C-rich ssDNAs were detected around 1.825 g/ml (Fig. 2F; fraction #1-4), indicating that they are single-stranded and nascent DNAs possibly originating from lagging daughter strands.

### MRE11 nuclease activity suppresses the generation of C-rich ssDNA

We investigated which nucleases are responsible for the generation of C-rich ssDNAs and C-circles. We first tested the impact of MRE11 nuclease activity inhibition in the generation of C-circles (Fig. 3A). We used Mirin, a small molecule inhibitor of MRE11 which inhibits MRE11 nuclease activity (30). Surprisingly, C-circles were increased in Mirin-treated condition in both U2OS and SaoS2 cells (Fig. 3B-C). Next, we performed a 4SET assay in Mirin-treated condition (Fig. 3D, S3A). Mirin treatment led to increases in C-rich ssDNAs mostly in the cytoplasm fractions, with C-rich ssDNAs not detectable in the nuclear fraction (Fig. 3D; G-rich probe).

**Figure 3.**
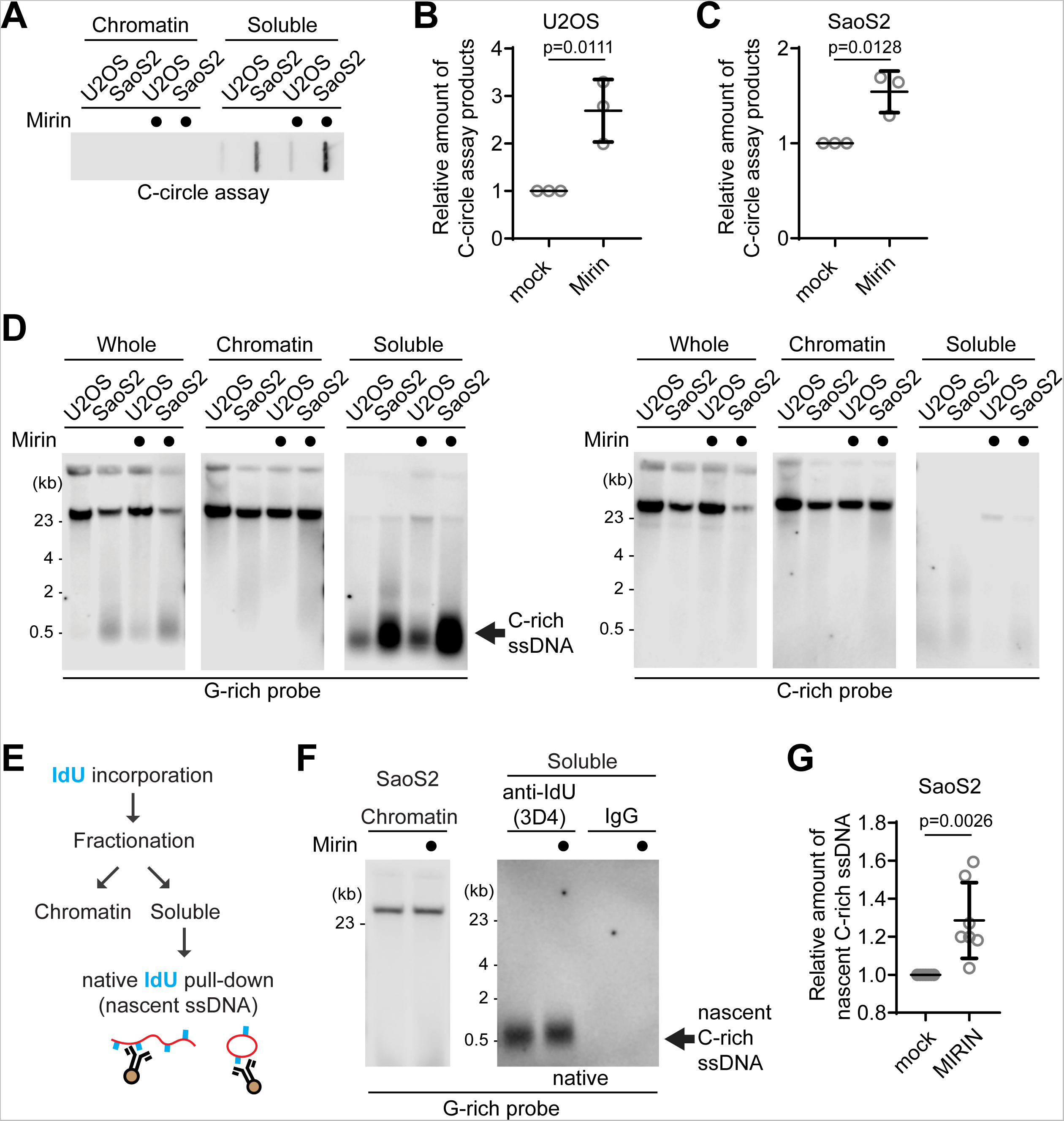
MRE11 nuclease inhibition leads to increases in C-circles and C-rich ssDNAs. **(A)** C-circle assay for U2OS and SaoS2 cells with or without Mirin (MRE11 nuclease inhibitor) treatment. Samples were fractionated into chromatin and soluble fractions. DNA amount: 100ng. U2OS and Saos2 cells were treated with 50 μM Mirin for 48 hr. **(B-C)** Quantification of the C-circle assay in A; U2OS cells (B), and SaoS2 cells (C), as the relative amount of C-circle assay products (mean ± SD; unpaired t-test). **(D)** 4SET assay for U2OS and SaoS2 cells with or without Mirin treatment after fractionation of samples into whole, chromatin, and soluble fractions. DNA amount: 600 ng. G-rich probe was used to detect C-rich telomeric sequences, and C-rich probe was used to detect G-rich telomeric sequences. C-rich single stranded DNAs are indicated by an arrow. **(E)** Illustration depicting the process of pulling down nascent single-stranded DNA from the soluble fraction incorporated with IdU. **(F)** Native IdU pulldown assay using IdU (3D4) antibody for SaoS2 cells treated with either Mirin (50 μM Mirin and 100 μM IdU for 20 hours) or mock (100 μM IdU for 20 hours). IgG-negative control; chromatin DNA-control for quantity. **(G)** Quantification of the native IdU-pulldown assay in F; Relative amount of nascent C-rich ssDNAs (mean ± SD; unpaired t-test).

We checked whether the impact of MRE11 inhibition in non-ALT cells to check whether Mirin-induced C-rich ssDNAs generation is ALT-specific phenotype. HeLa LT cells, telomerase-positive cells harboring long telomeres, were used (31). Mirin treatment in HeLa LT cells did not lead to generation of C-rich ssDNAs (Fig. S3B), indicating that C-rich ssDNAs are an ALT cancer-specific phenotype, as previously documented (24).

In addition to mirin, we repeated the experiment for the MRE11 nuclease inhibitors PFM01 and PFM39 which are derived from Mirin structure-based chemical screening (32). Both PFM01 and PFM39 treatments also increased C-rich ssDNAs (Fig. S3C), confirming that the impact of Mirin treatment is due to the inhibition of nuclease activity of MRE11. Additionally, MRE11 depletion led to a decrease in C-rich ssDNA accumulations in response to Mirin treatment, further confirming that Mirin’s effect is indeed attributed to its targeting of MRE11 nuclease activity (Fig. S3D-E).

To further validate that increased C-rich ssDNAs in Mirin treatment are nascently generated, we performed IdU pulldown in native condition of soluble DNAs to measure nascent C-rich ssDNA (Fig. 3E). Mirin treatment induces an increase in nascent C-rich ssDNA generation, indicating that MRE11 inhibition generates more C-rich ssDNAs which were nascently generated during DNA synthesis (Fig. 3F-G).

We subsequently investigated whether PML bodies contribute to the elevated C-rich ssDNAs observed upon Mirin treatment. In PML knockout SaoS2 cells, C-rich ssDNAs were less abundant and showed no increase upon Mirin treatment (Fig. S3F-G). Moreover, these knockout cells produced significantly fewer nascent C-rich ssDNAs, indicating that the increased C-rich ssDNAs upon Mirin treatment is primarily generated at APBs (Fig. S3H-I).

### C-rich ssDNAs are generated during Okazaki fragment processing

Prior research from both us and others has demonstrated that C-circles are produced during telomere replication and are predominantly present in the lagging strand compared to the leading strand (21, 22). Telomeres expose their G-rich ssDNAs during Okazaki fragment formation in the lagging strand (33). These G-rich ssDNAs can form higher order structures such as G-quadruplexes, which not only contribute to telomere fragility but also serve as substrates for nucleases, such as MRE11 and DNA2 (33–37).

We observed an increase in C-rich ssDNAs in conditions where MRE11 was inhibited (Fig. 4A), possibly preventing the processing of higher order structures in the lagging strands. Based on this observation, we hypothesized that C-rich ssDNAs are derived from lagging strand processing. To further investigate this idea, we decided to deplete DNA2, which is also involved in lagging strand processing at telomeres (37). We employed siRNA targeting DNA2 to deplete its levels in U2OS cells (Fig. S4A).

**Figure 4.**
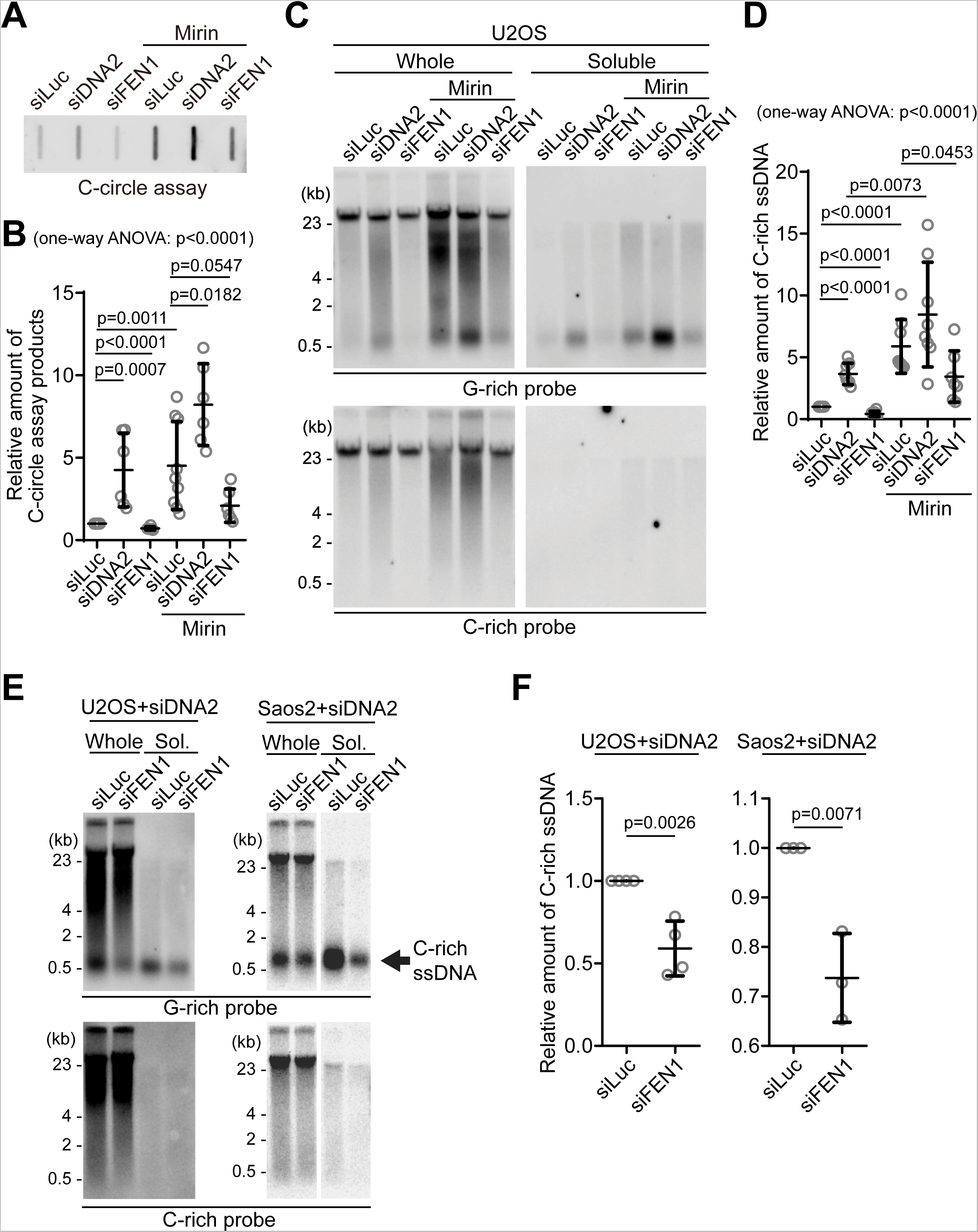
C-circles and C-rich ssDNAs are generated during Okazaki fragment processing. **(A)** C-circle assay for U2OS cells after transfection of siRNAs targeting DNA2 or FEN1 with or without Mirin treatment. DNA amount: 40 ng. **(B)** Quantification of the C-circle assay in A as the relative amount of C-circle assay products (mean ± SD; unpaired t-test). One-way ANOVA analysis; p<0.0001. **(C)** 4SET assay for U2OS cells after transfection of siRNAs targeting FEN1, DNA2, or control (siLuc) with or without Mirin treatment. Samples were fractionated into whole DNA or soluble fraction. DNA amount (whole: 210 ng, soluble: 70 ng). G-rich probe was used to detect C-rich telomeric sequences, and C-rich probe was used to detect G-rich telomeric sequences. **(D)** Quantification of the C-circle assay in C; as the relative amount of C-rich ssDNA (whole). (mean ± SD; unpaired t-test). One-way ANOVA analysis; p<0.0001. **(E)** 4SET assay for U2OS and SaoS2 cells after transfection of siRNAs targeting FEN1 and DNA2, or DNA2 and siLuc. DNA amount (whole: 210 ng, soluble: 70 ng). G-rich probe was used to detect C-rich telomeric sequences, and C-rich probe was used to detect G-rich telomeric sequences. C-rich single stranded DNAs are indicated by an arrow. **(F)** Quantification of the 4SET assay in E; as the relative amount of C-rich ssDNA (whole). (mean ± SD; unpaired t-test).

We initially measured C-circle levels in the DNA2 depletion condition (Fig. 4A) and found that, in line with previous reports, DNA2 depletion resulted in an increase in C-circle levels (Fig. 4B; siDNA2) (31). Interestingly, the combination of DNA2 depletion and Mirin treatment led to even higher C-circle levels, indicating an additive effect of inhibiting MRE11 and DNA2 depletion.

Then, we measure the C-rich ssDNAs in DNA2 depleted condition. We thus conducted a 4SET assay in the DNA2 depletion condition (Fig. 4C, S4B). DNA2 depletion alone caused an increase in C-rich ssDNA (Fig. 4C; siDNA2). Moreover, when Mirin was treated with DNA2 depletion, it further enhanced the accumulation of C-rich ssDNA (Fig. 4C; siDNA2 with Mirin). Therefore, in DNA2 depleted conditions, the increase in C-rich ssDNA is consistent with the increase in C-circles. However, DNA2 may not be the only nuclease involved in ssDNA generation.

As DNA2 has a role in removing the flap structures during Okazaki fragment processing, we tested if another flap nuclease, FEN1, might be involved in C-rich ssDNA generation (38). We depleted FEN1 using siRNA (Fig. S4C). Interestingly, FEN1 depletion led to a moderate decrease in C-circle levels (Fig. 4A, B; siFEN1). In the presence of Mirin treatment, FEN1 depletion again led to partial decrease in C-circle levels (Fig. 4A, B; siFEN1 with Mirin). Constantly, FEN1 depletion also led to a partial decrease in C-rich ssDNA generation (Fig. 4C-D, S4B) with or without Mirin treatment, which is completely opposite to our findings upon DNA2 depletion.

DNA2 exhibits a preference for cleaving long-flap structures from the 5’ end of the flap, whereas FEN1 is capable of cleaving the middle of the flap, as they are recruited by PCNA (Fig. S4D) (38). We hypothesize that FEN1 nuclease plays a role in generating C-rich ssDNAs by cutting the middle of 5’ flap structures when long-flap structures are not adequately processed by DNA2. To test this idea, we conducted a double-depletion experiment, simultaneously targeting FEN1 and DNA2, to determine if double depletion of FEN1 and DNA2 can impede the accumulation of C-rich ssDNAs in the DNA2 depleted condition (Fig. 4E-F, S4E). Double depletion of FEN1 and DNA2 exhibited decreased C-rich ssDNAs compared to single DNA2 depletion in both U2OS and SaoS2 cells, indicating that FEN1 partially contributes to the generation of C-rich ssDNAs, possibly cleaving the flap structures in the lagging strands.

Next, we explored whether the exposed ssDNAs in lagging strand telomeres themselves could contribute to an increase in C-rich ssDNAs. To test this idea, we utilized a PARP inhibitor, which is known to impact ssDNA generation (39, 40). We applied a modest dosage of Talazoparib, a PARP inhibitor, to induce ssDNA gaps (41). Remarkably, treatment with the PARP inhibitor resulted in an increase in C-rich ssDNAs, similar to what was observed with Mirin treatment (Fig. S4F), supporting the idea that C-rich ssDNAs are generated during DNA synthesis through ssDNA exposure and gap formation.

Next, we ruled out the possibility that C-rich ssDNA’s are generated from ssDNA or gaps from the leading strand. We focused on PRIMPOL, a Primase-polymerase known for its role in repriming and gap formations in leading strands (42–44). Notably, depletion of PRIMPOL did not show any changes in the levels of C-rich ssDNAs (Fig. S4G). Furthermore, we analyzed PRIMPOL knockout cells, which have a defect in gap formation on leading strands in response to DNA replication challenges (45). When we introduced either PRIMPOL wild-type (WT) or the phosphomimic and constitutively active mutant S255D into PRIMPOL knockout cells (46), we observed no changes in C-rich ssDNAs (Fig. S4H). These results also support the idea that C-rich ssDNAs are primarily derived from gap formation on the lagging strand rather than on the leading strand.

### ATRX suppresses the generation of C-rich ssDNAs

ATRX (α-thalassemia/mental retardation syndrome X-linked) is a tumor suppressor and histone chaperone. Mutations in the ATRX gene have been identified in many ALT cancers, or loss of ATRX expression is exhibited in a significant number of ALT positive cells (47, 48). Recent studies showed that reintroducing the ATRX gene expression using a doxycycline-inducible promoter, in U2OS cells, which lack ATRX due to homozygous deletion, effectively reversed the ALT phenotypes, such as C-circles, APBs, and accumulations of DNA/RNA hybrids (R-loops) at telomeres (49, 50). We used these U2OS^ATRX^ cells to test whether ATRX re-expression in U2OS^ATRX^ cells can affect the generation of C-rich ssDNAs. After 7 days of doxycycline treatment, U2OS^ATRX^ cells express ectopic ATRX (Fig. 5A).

**Figure 5.**
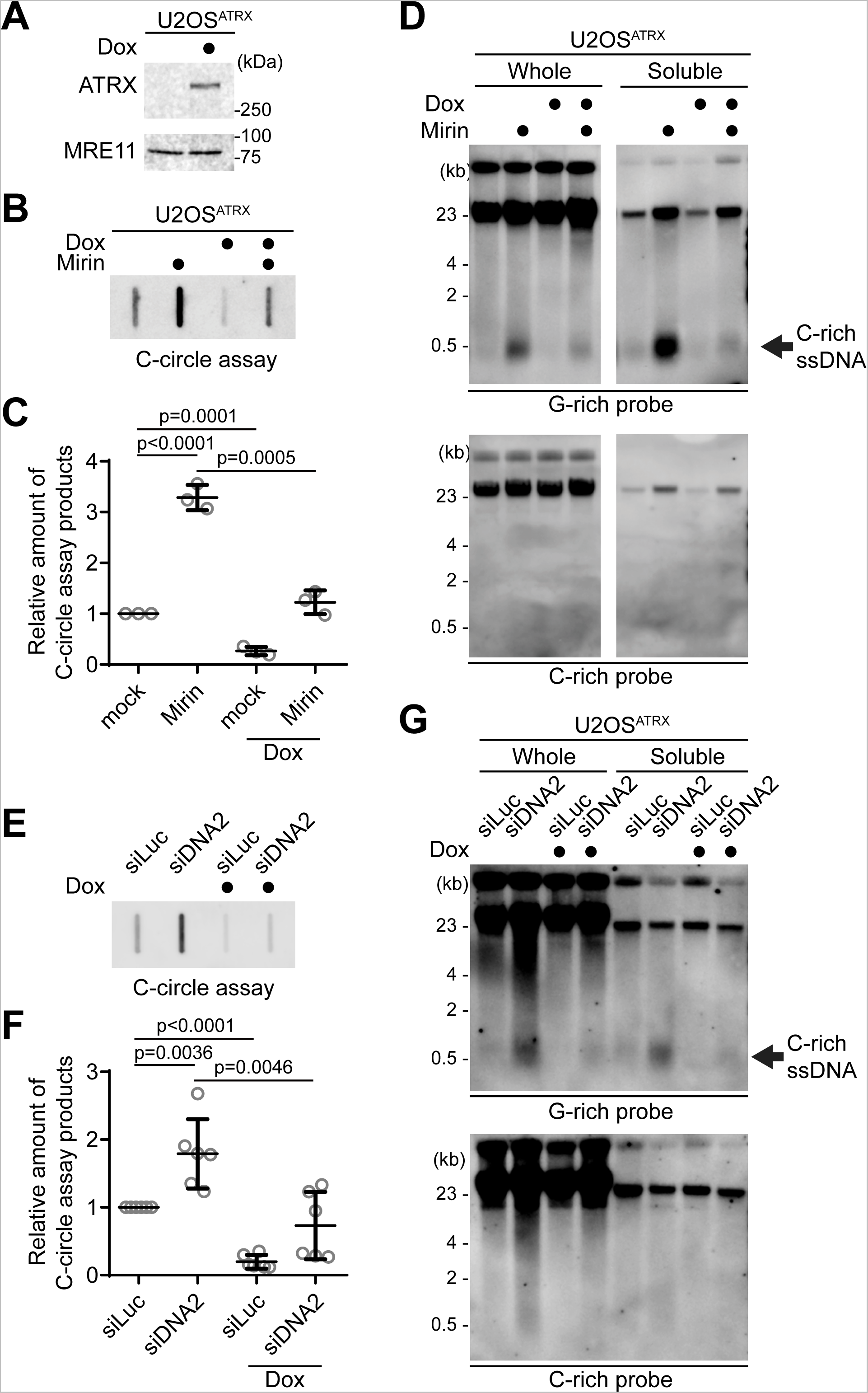
ATRX suppresses the C-rich ssDNA generation. **(A)** Western blot analysis of U2OS^ATRX^ cells with ectopic ATRX expression under a doxycycline-inducible promoter, using anti-ATRX and MRE11 antibodies for the validation of ATRX expression and MRE11 levels, respectively. **(B)** C-circle assay for U2OS^ATRX^ cells treated with doxycycline for one week or untreated, with or without Mirin treatment (50 μM Mirin for 48 hr). DNA amount: 10 ng. **(C)** Quantification of the C-circle assay in B as the relative amount of C-circle assay products (mean ± SD; unpaired t-test). **(D)** 4SET assay for U2OS^ATRX^ cells treated with doxycycline for one week or untreated, with or without Mirin treatment (50 μM Mirin for 48 hr). Samples were fractionated into soluble fraction or whole DNA. DNA amount (whole: 200 ng, soluble: 50 ng). G-rich probe was used to detect C-rich telomeric sequences, and C-rich probe was used to detect G-rich telomeric sequences. C-rich single stranded DNAs are indicated by an arrow. **(E)** C-circle assay for U2OS^ATRX^ cells treated with doxycycline for one week or left untreated. Cells were transfected with siRNAs targeting DNA2, or control (siLuc). DNA amount: 100 ng. **(F)** Quantification of the C-circle assay in B as the relative amount of C-circle assay products (mean ± SD; unpaired t-test). **(G)** 4SET assay for U2OS^ATRX^ cells treated with doxycycline for one week or untreated. Cells were transfected with siRNAs targeting DNA2, or control (siLuc). Samples were fractionated into soluble fraction or whole DNA. G-rich probe was used to detect C-rich telomeric sequences, and C-rich probe was used to detect G-rich telomeric sequences. C-rich single stranded DNAs are indicated by an arrow. (whole: 200 ng, soluble: 60 ng).

First, we measured C-circle levels in the doxycycline-induced ATRX expression condition (Fig. 5B). As expected, ATRX expression led to a decrease in C-circle levels. Interestingly, ATRX expression also counteracted Mirin treatment (Fig. 5B, C; Dox with Mirin), suggesting that the increase in C-circles induced by Mirin treatment are dependent on the absence of ATRX in U2OS cells.

Subsequently, we assessed telomeric C-rich ssDNAs under the condition of ATRX expression. ATRX expression in U2OS cells decreased the quantity of C-rich ssDNAs, and it alleviated the increase of C-rich ssDNAs induced by Mirin treatment (Fig. 5D, S5A). DNA2 depletion also led to an increase in C-circles in U2OS^ATRX^ cells (Fig. 5E, F; Dox with siDNA2). However, Doxycycline-induced ATRX expression diminished the change in both C-circles and telomeric C-rich ssDNA induced by DNA2 depletion (Fig. 5G, S5B). These results indicate that loss of ATRX is a crucial factor in the generation of C-rich ssDNAs in U2OS cells.

Considering that ATRX is involved in resolving G-quadruplexes (G4) at telomeres and its depletion can lead to the accumulation of R-loop structures at telomeres associated with G4 (49, 51), we predicted that DNA/RNA hybrid formation in the leading strand could produce C-rich ssDNAs by causing gaps in the unhybridized strand in the lagging strand. We found that FANCM depletion (Fig. S5C), which induces R-loop accumulation at ALT telomeres (Fig. S5D) (52, 53), led to an increase in C-rich ssDNAs (Fig. S5E), which is consistent with the previous observation (54). To further validate this idea, we analyzed RAD51AP1 KO cells (Fig. S5F) (55), which are known to exhibit reduced G-quadruplex-associated R-loop formations at ALT telomeres (56–58). RAD51AP1 KO cells displayed lower levels of C-rich ssDNAs compared to Control cells (Fig. S5G). Collectively, these results indicate that loss of ATRX is a crucial factor in the generation of C-rich ssDNAs, likely facilitated by the accumulation of R-loops at telomeres.

Nevertheless, previous studies demonstrate that the loss of ATRX genes is insufficient to induce the ALT phenotype (47, 59–61), We hypothesized that C-rich ssDNA are abundant in ALT positive cells, so we confirmed that ATRX loss alone would not generate ssDNA. As expected, when we introduced shRNA targeting the ATRX gene in HeLa LT cells, and we did not observe the generation of C-rich ssDNAs in HeLa LT cells after ATRX depletion, even when subjected to Mirin or PARPi treatment in our 24-hour treatment experiment (Fig. S5H). These results indicate that ATRX may not be the sole determinant in the regulation of C-rich ssDNA formation in HeLa LT cells under the specific short-term experimental conditions we employed.

### CST complex-mediated priming and subsequent strand displacements by DNA polymerase delta generates C-rich ssDNAs

The CST (CTC1/STN1/TEN1)-PP (Primase/DNA polymerase alpha) complex plays a crucial role in the C-strand fill-in processes at telomeres (62). It recognizes exposed G-rich strands, including higher-order structures like as G-quadruplex, on the parental lagging strand during telomere replication and initiates the filling in of the complementary C-strand, generating additional Okazaki fragments that serve to remove the gaps in the lagging strand and ensure the telomere replication (63–67). Because the exposed ssDNAs in lagging strand telomeres are priming sites for CST complex, we hypothesized that C-rich ssDNAs could be generated by additional Okazaki fragments formation by CST complex-mediated DNA priming (Fig. 6A). We depleted STN1, a component of CST complex, using shRNA targeting STN1 (67) (Fig. S6A). In line with the previous report, depletion of STN1 led to a reduction of C-circles in Mirin-treated U2OS cells (Fig. 6B, C). We performed a 4SET assay to measure the levels of C-rich ssDNAs in conditions where STN1 was depleted. Surprisingly, the depletion of STN1 led to a striking reduction in C-rich ssDNAs in both U2OS and SaOS2 cells (Fig. 6D-E, S6B-C). Furthermore, STN1 depletion also resulted in the dramatic abolishment of C-rich ssDNAs in response to Mirin treatment. These findings suggest that STN1 plays a crucial role in the generation of C-rich ssDNAs and highlights its involvement in the response to Mirin treatment in these cells.

**Figure 6.**
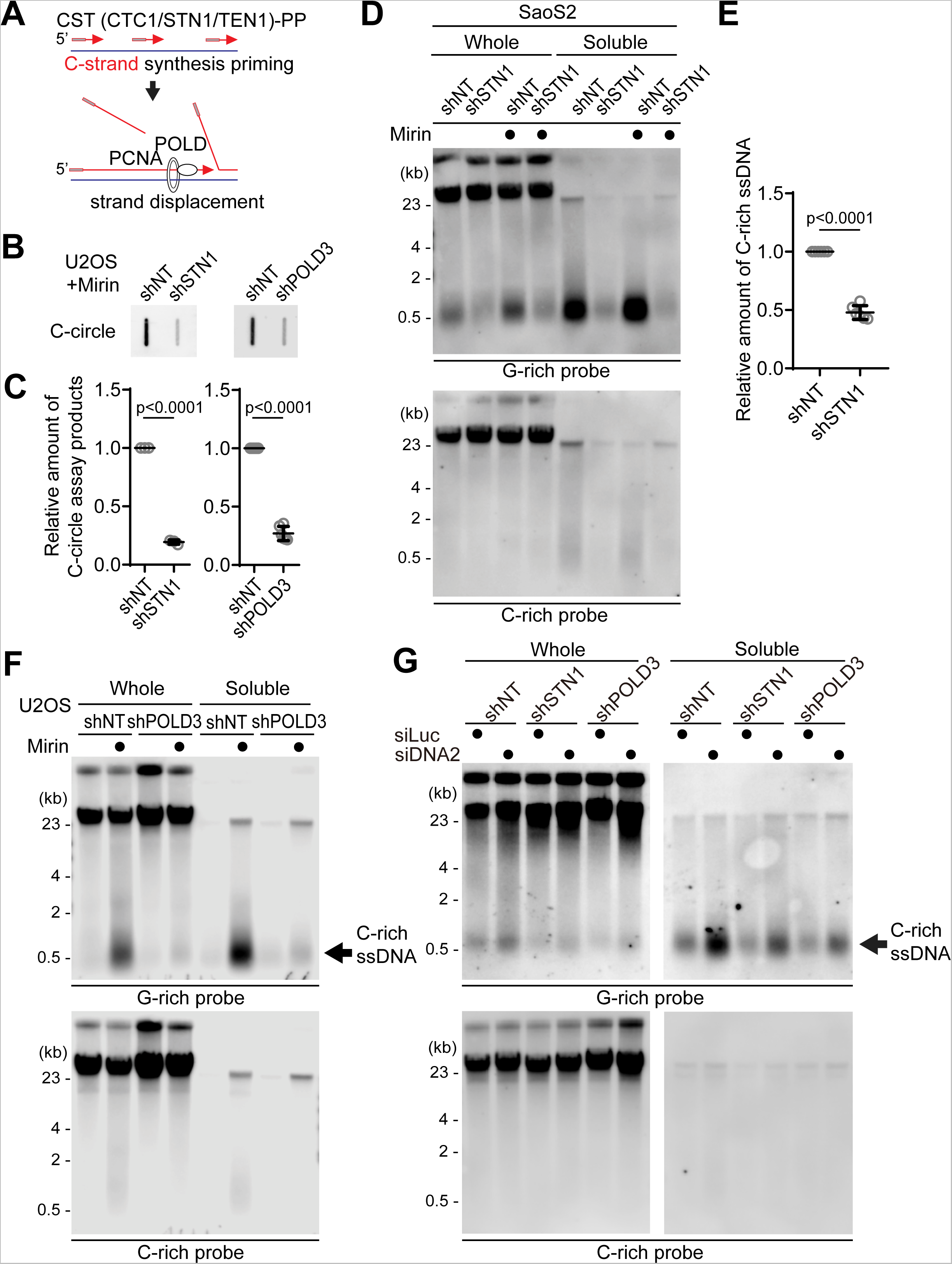
C-circles and C-rich ssDNAs are generated through C-strand priming and subsequent strand displacement. **(A)** Illustration depicting C-strand synthesis priming by CST-PP (CTC1/STN1/TEN1-Primase-Polymerase alpha)-complex and the strand displacement of Okazaki fragments by PCNA and DNA polymerase delta (POLD) during DNA replication on the lagging strand. **(B)** C-circle assay for U2OS cells expressing shRNAs targeting STN1, POLD3, or non-targeting (NT) control. Cells were treated with Mirin (50 μM, 48 hr) DNA amount: left - 100ng, right - 30ng. **(C)** Quantification of the C-circle assay in B as the relative amount of C-circle assay products (mean ± SD; unpaired t-test). **(D)** 4SET assay for SaoS2 expressing shRNAs targeting STN1, or non-targeting (NT) control. Cells were treated with Mirin (50 μM, 48 hr). Samples were fractionated into soluble fraction or whole DNA. G-rich probe was used to detect C-rich telomeric sequences, and C-rich probe was used to detect G-rich telomeric sequences. C-rich single stranded DNAs are indicated by an arrow. DNA amount (Saos2 whole: 200 ng, soluble: 60 ng). **(E)** Quantification of the 4SET assay in D; as the relative amount of C-rich ssDNA (whole). (mean ± SD; unpaired t-test). **(F)** 4SET assay for U2OS cells expressing shRNAs targeting POLD3, or non-targeting (NT) control. Samples were fractionated into soluble fraction or whole DNA. G-rich probe was used to detect C-rich telomeric sequences, and C-rich probe was used to detect G-rich telomeric sequences. C-rich single stranded DNAs are indicated by an arrow. DNA amount (whole: 300 ng, soluble: 100 ng). **(G)** 4SET assay for U2OS cells expressing shRNAs targeting STN1, POLD3, or non-targeting (NT) control after transfection of siRNAs targeting DNA2 or control (siLuc). Samples were fractionated into soluble fraction or whole DNA. G-rich probe was used to detect C-rich telomeric sequences, and C-rich probe was used to detect G-rich telomeric sequences. C-rich single stranded DNAs are indicated by an arrow. DNA amount (whole: 210 ng, soluble: 70 ng).

Primed Okazaki fragments, created by DNA polymerase alpha, are subsequently extended by DNA polymerase delta (38). This sequential process ensures the continuous synthesis of the lagging strand during DNA replication and creates a flap structure as polymerase delta displaces the downstream DNA strand. Particularly in ALT cells, the excessive strand displacement activity of polymerase delta, working with the BLM helicase and other associated factors, plays a crucial role in ALT telomere elongation through BIR (5, 68–70). We hypothesized that the primed C-strand Okazaki fragments, initiated by the CST-PP complex, undergo prolonged extension by DNA polymerase delta and BLM helicase, creating a flap structure. This unique excessive strand displacement activity in ALT cells allows for the complete displacement of downstream DNA, resulting in the accumulation of C-rich ssDNA (Fig. 6A).

To test this idea, we depleted POLD3, a component of DNA polymerase delta, using shRNA targeting POLD3 (71, 72). Depletion of POLD3 led to a decrease in C-circles in Mirin treated U2OS cells. We conducted a 4SET assay to measure the C-rich ssDNAs in POLD3-depleted cells. Strikingly, POLD3-depletion reduced the C-rich ssDNAs in U2OS cells (Fig. 6F, S6D). Moreover, POLD3-depletion also abolished the C-rich ssDNA in response to Mirin treatment.

To further validate the involvement of DNA polymerase alpha and delta in the generation of C-rich ssDNAs, we conducted 4SET assay after treating low dosages of Aphidicolin (a DNA polymerase alpha/delta inhibitor) and CD437 (a DNA polymerase alpha inhibitor) to partially inhibit their actions. The results were in line with our hypothesis, as both Aphidicolin and CD437 treatments led to decreases in C-rich ssDNAs (Fig. S6E), confirming that C-rich ssDNAs are indeed derived from lagging strand processes mediated by DNA polymerase alpha and delta. The inhibition of these DNA polymerases resulted in reduced C-rich ssDNA formation, supporting their role in this process during telomere replication.

As we hypothesized, we observed that BLM knockout cells showed reduced levels of C-rich ssDNAs (Fig. S6F-G). These results support that BLM helicase is involved in the generation of C-rich ssDNAs in the context of ALT telomere replication through BIR. The reduced levels of C-rich ssDNAs upon BLM depletion suggest that BLM’s excessive strand displacement activity likely plays a crucial role in this process.

Furthermore, STN1- and POLD3-depletion also mitigated the increase of C-rich ssDNAs induced by DNA2 depletion (Fig. 6G, S6H), supporting the idea that C-strand synthesis priming followed by excessive strand displacement is required for the generation of C-rich ssDNAs in ALT cells. Finally, we assessed nascent C-rich ssDNA under STN1- depleted conditions (Fig. 7A). The depletion of STN1 resulted in reduced nascent C-rich ssDNA (Fig. 7B), again demonstrating that C-rich ssDNAs are nascently generated and arise from CST-complex mediated priming at the lagging strands.

**Figure 7.**
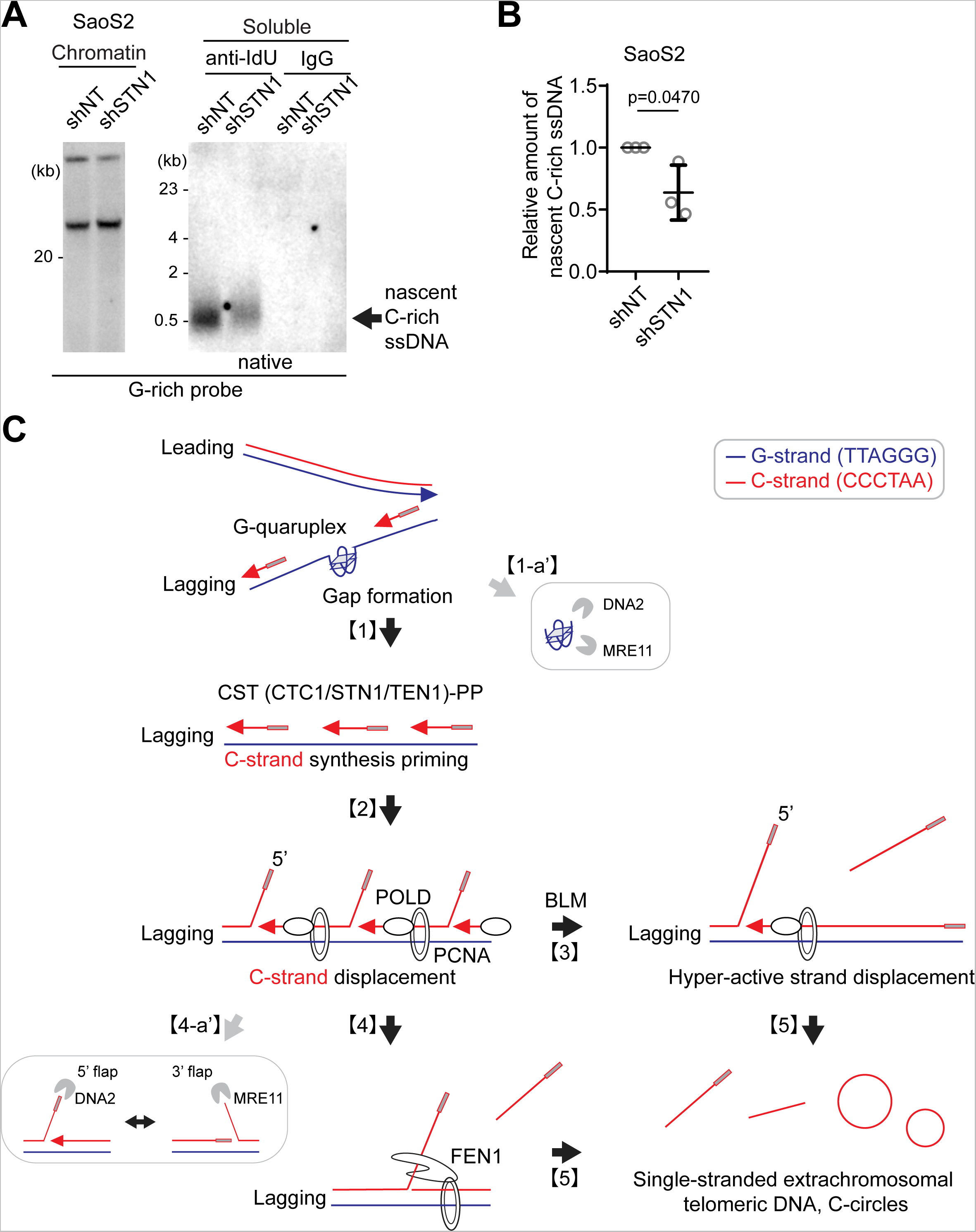
Origins of C-circles and their precursor linear C-rich single-stranded DNAs: Excessive strand displacements from lagging strand telomeres. **(A)** Native IdU pulldown was conducted using IdU (3D4) antibody for SaoS2 cells expressing shRNAs targeting STN1, or non-targeting (NT) control. IgG served as a negative control. Chromatin DNA was also included to control for DNA quantity. **(B)** Quantification of the native IdU-pulldown assay in F; Relative amount of nascent C-rich ssDNAs (mean ± SD; unpaired t-test). **(C) Working model:** Accumulations of G-quadruplex (G4) structures and G4-associated R-loops at telomeres may contribute to the generation of C-rich single-stranded DNAs (ssDNAs) by potentially exposing parental lagging strands, which can be recognized by the CST-PP (CTC1-STN1-TEN1-Polα-Primase) complex **[1]**. The CST-PP complex exhibits priming activity in the exposed gaps or G4 structures in lagging strands, which are subsequently extended by DNA polymerase delta **[2]**. As a result, primed C-strand Okazaki fragments are initiated by the CST-PP complex and undergo prolonged extension facilitated by DNA polymerase delta and BLM helicase, leading to the formation of long flap structures **[3]**. In the context where long flap structures are not adequately processed by DNA2, the FEN1 nuclease may contribute to the generation of C-rich ssDNAs by cleaving the middle of 5’ flap structures **[4]**. Notably, the unique excessive strand displacement activity observed in ALT cells enables the complete displacement of downstream DNA, thereby promoting the accumulation of C-rich single-stranded DNA **[5]**. DNA2 and MRE11 are potentially involved in lagging strand telomere processes by participating in the removal of G4 structures **[1-a’]** and processing flap structures in Okazaki fragments on the lagging strands **[4-a’]**.

## Discussion

### 4SET, a sensitive method to detect extrachromosomal single stranded DNAs

We demonstrate a highly sensitive and accessible method to detect extrachromosomal telomeric ssDNAs called 4SET (Strand-Specific Southern-blot for Single-stranded Extrachromosomal Telomeres) assay. This technique can be performed within two days by molecular biologists at an intermediate level of expertise, without the need for specialized or expensive laboratory equipment, allowing us to investigate single-stranded extrachromosomal telomeres in a time-efficient and cost-effective manner. Additionally, this assay offers a strand-specific approach, allowing for the discrimination of specific telomere strands during analysis. For example, by employing the 4SET assay, here we proposed the model of underlying mechanisms of generation of single-stranded extrachromosomal telomeres which are derived from strand displacement of lagging strands. Furthermore, the simplicity and efficiency of the 4SET assay make it suitable for implementation in various research settings, such as assessing other repeat sequences of interest.

### Generation of C-rich single stranded DNAs are suppressed by MRE11, DNA2 nucleases

In our study, we explored the roles of MRE11 and DNA2 nucleases in the generation of C-rich single-stranded DNAs. MRE11, along with RAD50 and NBS1 forms MRN complex, is involved in DNA repair and replication processes through its nuclease activity which initiates the resection of DNA ends in DNA double-stranded breaks (DSBs) for homology-directed recombination (HDR) (73). In addition to its role at DSBs, MRN associates with unperturbed replication fork as well as stalled replication forks (74–77). In particular, MRN complex is localized at telomeres during S phase through the NBS1-TRF2 interaction (78, 79). Moreover, MRN localizes to ALT-associated PML bodies (APBs) in ALT cells and is essential for ALT activity and the generation of T-circles, as shown by the observation that depletion of MRN using shRNA leads to a decrease in APBs and compromises these processes (80–82). However, depletion of MRE11 itself does not affect the generation of C-circles (31), although sequestration of the MRN complex by overexpressing the Sp100 protein or the ATRX protein dramatically reduces the C-circle levels (16, 49, 83, 84). Further analysis is needed to clarify the specific mechanisms underlying these observations and their implications for telomere maintenance in ALT cells.

Here, our study demonstrated that inhibiting the nuclease activity of MRE11 results in an increase in the generation of C-rich ssDNAs and C-circles. These findings suggest that the nuclease activity of MRE11 plays a role in regulating the levels of C-rich ssDNAs and C-circles, potentially contributing to the resolution of higher-order structures, such as G-quadruplexes in lagging strands (Fig. 7C; [1-a’]) or 5’ flap structures in Okazaki fragments (Fig. 7C; [4-a’]). Notably, yeast genetics showed that *mre11* and the mre11 nuclease mutant are synthetic lethal with *rad27* (yeast homolog of human FEN1) (85), even though in vitro biochemistry experiments did not reveal preferential MRE11 nuclease activity for flap structures (86).

Additionally, our study demonstrated that depletion of DNA2 leads to an increase in the generation of C-rich ssDNAs and C-circles, consistent with previous reports that showed elevated C-circles and APBs upon DNA2 depletion (31, 38, 87). DNA2 plays a crucial role in resolving G-quadruplex structures in lagging strand telomeres, which likely contributes to the observed accumulation of C-rich ssDNAs upon DNA2 depletion (Fig. 7C; [1-a’]).

DNA2 also plays a crucial role in Okazaki fragment processing, particularly in the removal of flap structures (38, 88). Recent structural studies have unveiled that DNA2 adopts a cylindrical shape with a tunnel through which ssDNA passes, and its nuclease domain is embedded within this tunnel (89). DNA2 efficiently trims ssDNA from the ssDNA end and preferentially processes long flaps (Fig. 7C; [4-a’]). This process relies on its interaction with RPA (Replication Protein A) (89). We propose that DNA2 depletion results in an increased presence of long flaps, which are partly processed by FEN1 or engaged in strand displacement by DNA polymerase delta. FEN1 nuclease may contribute to the generation of C-rich ssDNAs by cleaving the middle of 5’ flap structures when long-flap structures are not adequately processed by DNA2 (Fig. 7C; [4]). However, the contribution of FEN1-mediated cleavage in C-rich ssDNAs may not be substantial when the flaps are long and bound by RPA. Indeed, further study is essential to fully elucidate the precise mechanisms by which these enzymes coordinate their activities for Okazaki fragment processes in the lagging strand, particularly at ALT telomeres.

### C-rich ssDNAs are generated by strand displacement of lagging strands

Our results indicate that the loss of ATRX is a crucial factor in the generation of C-rich ssDNAs, likely facilitated by the accumulations of G-quadruplex (G4) structures and G4-associated R-loops at telomeres. This is supported by FANCM depletion and RAD51AP1 KO data showing that R-loop formation may contribute to the generation of C-rich ssDNAs, possibly by exposing parental lagging strands that can be recognized by the CST-PP complex (Fig. 7C; [1]).

The CST-PP complex has critical roles during telomere replication in particular of the lagging strand (67). Previous reports demonstrated that CST complex has a role in resolving G4 structures at telomeres as well as genome wide (90). Interestingly, CST complex is detected in APBs (66). Depletion of CST leads to replication defects in ALT cells, accompanied by decreases in T-circles and C-circles (66). We interpret that CST’s priming activity in exposed gap or G4 structures in lagging strands is responsible for the generation of C-rich ssDNAs and C-circles. These primed Okazaki fragments, created by DNA polymerase alpha, are subsequently extended by DNA polymerase delta (Fig. 7C; [2]).

In ALT cells, the interplay between polymerase delta’s excessive strand displacement activity, along with BLM helicase and other associated factors, plays a crucial role in ALT telomere elongation through Break-Induced Replication. We propose that primed C-strand Okazaki fragments, initiated by the CST-PP (CTC1-STN1-TEN1-Polα-Primase) complex, undergo prolonged extension facilitated by the DNA polymerase delta and the BLM helicase, resulting in the formation of long flap structures (Fig. 7C; [3]). The unique excessive strand displacement activity in ALT cells enables the complete displacement of downstream DNA, leading to the accumulation of C-rich single-stranded DNA (Fig. 7C; [5]).

Further investigations into the mechanisms underlying this process could provide valuable insights into the complex interplay between ATRX, G-quadruplex structures, R-loops, CST, and BLM/POLD3, contributing to our understanding of ALT telomere maintenance.

## Materials and Methods

### Reagents

Mirin (Selleckchem, SML1839; Sigma-aldrich, M9948), PFM01 (Sigma-Aldrich, SML1735), PFM39(Sigma-Aldrich, SML1839), Talazoparib (Selleckchem, S7048), aphidicolin (Sigma-Aldrich, A0781), CD437 (Sigma-Aldrich, C5865), Agarose (Sigma-Aldrich, A9539), SSC buffer (Sigma-Aldrich, S6639), CDP-star (Roche, 11685627001), Blocking reagent (Roche, 11096176001), PBS (Genesee, 25-507B), Glycogen (GlycoBlue Coprecipitant, Invitrogen, AM9515), anti-beta actin (Sigma-Aldrich, A5316), anti-Histone H3 (Cell signaling, 2650), anti-ATRX (Santa Cruz, sc-15408), anti-hMRE-11 (Calbiochem, PC388), anti-GAPDH (Cell signaling, 97166), TB Green Premix Ex Taq II (Tli RNase H Plus) (Takara, #RR820A), RevertAid First Strand cDNA synthesis kit (Thermo scientific, K1621), Lipofectamine 3000 (Thermo scientific, L3000001), TransIT (Mirus, MIR2300), Qubit dsDNA HS Assay Kit (Invitrogen, Q32851), Nylon Membranes-positively charged, (Roche, 11417240001), exo^-^ Klenow fragment (NEB, M0212), T4 DNA polymerase (NEB, M0203), Lambda exonuclease (NEB, M0262), DIG-dUTP (biorbyt, orb533128), NxGen phi29 (Biosearch Technologies, 30221), iododeoxyuridine (Sigma-Aldrich, I7125).

### Cell culture

U2OS, SaoS2, and HeLa LT cells were maintained in Dulbecco’s Modified Eagle Medium (DMEM) supplemented with 10% fetal bovine serum, Penicillin-Streptomycin (Gibco, 15070063). To prevent mycoplasma contaminations, we regularly used Plasmocin Prophylactic (InvivoGen) to maintain the cells according to the manufacturer’s protocol.

### siRNA-mediated knockdown

siRNA oligos were purchased from Sigma-Aldrich (10 nmol) and purified by desalting. The dTdT overhangs were added to ensure the stability of siRNA. The siRNAs were transfected into the cells at a final concentration of 20 nM, along with 2 μl of Lipofectamine 3000 (Thermo scientific, L3000001) according to the manufacturer’s protocol.

The following siRNAs were used:

siLuc: UCGAAGUAUUCCGCGUACGUU,

siDNA2#1: CAGUAUCUCCUCUAGCUAGUU (91),

siDNA2#2: AUAGCCAGUAGUAUUCGAUUU (91),

siFEN1#1: UGACAUCAAGAGCUACUUU (92),

siFEN#2: CCAUUCGCAUGAUGGAGAA (92),

siPRIMPOL: GAGGAAACCGUUGUCCUCAGUG (43),

siFANCM: AAGCUCAUAAAGCUCUCGGAA (52),

siMRE11: ACAGGAGAAGAGAUCAACU (93),

### Lentiviral shRNA knockdown and sgRNA knockout

Lentiviruses were generated by transfecting shRNA vectors targeting human POLD3 (71), STN1 (67), ATRX(47), or the control vector, or sgRNA vectors targeting human PML exon 3 (sense: CACCGGcggtaccagcgcgactacg, antisense: AAACcgtagtcgcgctggtaccgCC) inserted in pLentiV2 vector (Addgene, 52961) along with pMD2.G and psPAX into 293T cells. Transfection was carried out using TransIT (Mirus) following the manufacturer’s protocol. After 24 hours, the medium was replaced with fresh medium, and cells were incubated for an additional 24 hours. Supernatants containing the lentiviruses were collected and passed through a 0.45 μm syringe filter. The collected supernatants were then concentrated using 4x virus concentrator solution. This solution consisted of phosphate-buffered saline (PBS) at pH 7.2 containing 1.2 M sodium chloride (NaCl) and 40% (v/w) PEG-8000. The concentrated virus pellets were used to infect target cells, and 0.5 μg/ml polybrene was added to enhance viral transduction. Cells were selected under either 5 μg/ml Blasticidin-S (Invivogen) for two weeks or 5 μg/ml Puromycin (Invivogen) for 5 days to establish stable cell lines expressing the desired shRNA or sgRNA. After selection, cells are cloned and confirmed by RT-qPCR or Western blot. Proper biosafety procedures were followed when working with lentiviruses.

### Plasmid

cDNA encoding PRIMPOL was synthesized from Integrated DNA Technologies (IDT) and cloned into pCS2(+) NLS-mCherry vector. PRIMPOL S255D mutant was generated using Q5 site-directed mutagenesis kit (NEB). The pCS2 NLS-mCherry-PRIMPOL WT or S255D vectors were used for transfection in U2OS PRIMPOL KO cells using TransIT (Mirus) according to the manufacturer’s protocol. cDNA expression and transfection efficiency were assessed by mCherry signal.

### Strand-Specific Southern for Single-stranded Extrachromosomal Telomeres (4SET)

#### DNA preparation procedure

Cells were harvested using Trypsin-EDTA and then divided into separate tubes for fractionation. The cells were pelleted via centrifugation (500 g, 5 min) and washed with PBS. Cell pellets can be preserved at −80°C prior to 4SET analysis. For the whole DNA fraction, cells were lysed by adding lysis buffer (0.1 M Tris pH 8.0, 0.1 M EDTA, 1% SDS). To obtain the chromatin and soluble fractions, cells were treated with extraction buffer (PBS, 5 mM EDTA, 0.1% Triton X-100) and subjected to pipetting over ten times. The cell lysate was subsequently centrifuged at 1500 g for 5 min to separate the chromatin pellet from the soluble supernatant. The chromatin pellet was then lysed using lysis buffer. After obtaining the whole, chromatin, and soluble fractions, NaCl was added to each fraction (final concentration 1.6 M NaCl). The fractions were then centrifuged at 4°C, 15,000 g for 20 minutes to remove cell debris and obtain the DNA-containing supernatant. To precipitate the DNA, Glycogen (GlycoBlue Coprecipitant) was added to the supernatant, followed by the addition of 75% of the supernatant volume of isopropanol. The mixture was subjected to additional centrifugation at 4°C, 15,000 g for 20 minutes to obtain the DNA pellet. Subsequently, the DNA pellet was washed twice with 70% ethanol by centrifuging at 15,000 g for 5 minutes each time. Finally, the DNA pellet was dissolved in DW (distilled water) and incubated overnight at 4 °C to ensure complete dissolution. The DNA concentration was measured using Qubit (Qubit dsDNA HS Assay Kit, Invitrogen) or a nanodrop spectrophotometer.

#### Southern blot procedure

For electrophoresis, 210 ng of whole DNA, measured using Qubit, was loaded into 0.6% Agarose gel and run at 40 V (2.9V/cm) in TAE (Tris-acetate EDTA) buffer. Subsequently, the agarose gel underwent sequential incubations in the following solutions: Depurination solution (0.2 M HCl) for 5 minutes, Denaturation solution (0.5 M NaOH, 1.5 M NaCl) for 5 minutes, and Neutralization Buffer (0.5 M Tris pH 8.0, 1.5 M NaCl) for 20 minutes. The DNA was then transferred onto a Nylon membrane (Nylon Membranes, positively charged, Roche) using a Vacuum Blotting System (VacuGene XL) at 30 mbar for 1 hour in 4xSSC (Saline Sodium Citrate) buffer. Following this, the membrane was incubated with 1 nM DIG (Digoxigenin)-labeled Telomere probes in Hybridization solution (DIG Easy Hyb, Roche) at 42°C. Afterward, the membrane underwent two washes with Wash buffer 1 (2xSSC, 0.1% SDS) and two washes with Wash buffer 2 (0.5xSSC, 0.1% SDS) for 10 minutes each. The membrane was then blocked with Blocking solution (1% Blocking reagent, Roche 11096176001, in Maleic acid buffer pH 7.4) for 30 minutes, followed by incubation with 0.2% Anti-Digoxigenin-AP Fab fragments (Roche) in Blocking solution for 1 hour. Subsequently, the membrane underwent two washes with Wash buffer 3 (Maleic acid, 0.3% tween-20) for 15 minutes each. Finally, the membrane was incubated with 1% CDP-star (Roche 11759051001) in AP buffer (50 mM Tris pH 9.5, 100 mM NaCl) for detection, which was performed using the Odyssey Imaging System (LI-COR 2800).

#### Strand specific probe generation procedure

For the generation of strand-specific probes, we made minor modifications to the previous protocol (28). Annealed templates (TC, TG, and UP* in the list below) were generated at a concentration of 40 μM in NEBuffer#2.1 (NEB) buffer. The templates were then incubated with 50 units of exo-Klenow enzyme (NEB) and 1 mM dNTP with 0.35 mM DIG-dUTP (biorbyt) at 25°C overnight, followed by heat inactivation at 75°C for 20 minutes. Subsequently, we performed a 10-unit T4 DNA polymerase reaction to improve specificity. The double-stranded oligos were purified using NucleoSpin Gel and PCR clean-up (MN) with NTI buffer following the manufacturer’s instructions. Next, a 20-unit lambda exonuclease reaction was carried out to obtain single-stranded oligos for 1 hour at 37°C. Finally, the single-stranded telomere probes were purified using NucleoSpin Gel and PCR clean-up (MN) with NTC buffer.

Following oligos were used:

Telomere C-rich probe template (TC):

5’[Phos]GGGTTAGGGTTAGGGTTAGGGTTAGGGTTAGGGTTAGGGTTAGATAGTTGA GAGTC-3’

Telomere G-rich probe template (TG):

5’[Phos]CCCTAACCCTAACCCTAACCCTAACCCTAACCCTAACCCTAGATAGTTGAGA GTC-3’

Universal priming oligo (UP*): 5’-[Phos]GACTCTCAAC*T*A*T*C*T*A-3’

(*) indicates phosphorothioate bond

### CsCl separation (Cesium gradient analysis)

CsCl separation was performed as previously with minor modification (21). Briefly, cells were treated with 100 μM IdU for 20 hours, in conjugation with 10 nM FudR and 100 nM PARG inhibitor to promote IdU-incorporation and the recovery of C-rich ssDNAs. Post-incorporation, genomic DNA was extracted, yielding approximately 200 μg using the Puregene Core Kit A from Qiagen. The extracted DNA was then subjected to digestion using a combination of HinfI and RsaI enzymes. Following digestion, the DNA samples were mixed with a CsCl solution to achieve a target density of 1.770 g/ml. Centrifugation of these samples was performed at a speed of 55,000 rpm for 20 hours, utilizing a VTi-90 vertical rotor from Beckman. After centrifugation, the fractions were collected and a slot blot analysis for each fraction was executed.

### Cell fractionation using E1 and E2 buffer

Cells were lysed in E1 buffer (140 mM NaCl, 1 mM EDTA, 50 mM HEPES-KOH pH 7.5, 0.5% NP-40, 10% glycerol, and 0.25% Triton X-100). After mixing, the mixture was centrifuged at 2000 g for 2 minutes. The supernatant was prepared as the cytoplasm fraction. The pellet was then washed twice with E1 buffer, centrifuging at 1000 g for 2 minutes each time. On the second wash, the pellet was incubated on ice for 15 minutes. Subsequently, the pellet was resuspended in E2 buffer (200 mM NaCl, 1 mM EDTA, 10 mM Tris pH 8, and 0.5 mM EGTA pH 8). After a 20-minute incubation on ice, the supernatant was collected (nucleoplasm fraction). The remaining pellet was designated as the chromatin fraction.

### Nascent C-rich ssDNA pull-down (native IdU-immunoprecipitation)

IdU-pulldown was conducted under native conditions (without heat or denaturation), as detailed described (37). Cells were exposed to 100 μM IdU for 20 hours. Genomic DNAs from chromatin and soluble fractions were harvested from approximately 10 million cells. 1 µg of this genomic DNA from soluble fraction was mixed with 3 µg of anti-IdU/BrdU antibody (3D4 clone, BD Science) in BrdU-IP buffer composed of PBS and 0.0625% Triton X-100 overnight. This mixture was then subjected to a 1-hour incubation with Protein G magnetic beads. The immunoprecipitated DNA was subsequently eluted using an elution buffer (10 mM Tris pH 8 and 1% SDS). The elution was performed at room temperature and agitated at 1000 rpm for 1 hour. This was then purified using NucleoSpin Gel and PCR clean-up (MN) with NTC buffer. Eluted samples were analyzed using a native agarose gel electrophoresis.

### C-circle assay

Genomic DNAs were subjected to incubation with 10 units of phi29 polymerase (NxGen phi29), 0.2 mM dNTP mix, and 1X phi29 buffer at 30°C for 12 hours, followed by a 20-minute incubation at 65°C. Subsequently, the samples were loaded onto a Nylon membrane using a slot-blot technique. The subsequent steps of hybridization and detection followed the same procedure as described in the 4SET method.

### Western Blot

The cell pellet, nucleus pellet, and cytoplasm supernatant were treated with Laemmli loading buffer (2% SDS, 5% beta-mercaptoethanol, 10% Glycerol, 0.002% Bromophenol Blue, 62.5 mM Tris-base) and subjected to 10 seconds of ultrasonication. The prepared samples were then separated on a Bis-Tris 4-12% polyacrylamide gel and transferred to a PVDF membrane using the iBlot 2 Dry Blotting System. After blocking with 5% skim milk in PBS-T (PBS-0.1% Tween-20) for 1 hour at room temperature, the membrane was incubated with the primary antibody overnight at 4°C. Following PBS-T wash, the membrane was incubated with the secondary antibody for 1 hour at room temperature. After an additional PBS-T wash, the ECL solution was applied, and detection was performed using the Odyssey Imaging System (LI-COR 2800).

### Antibodies

The following antibodies were used for Western Blot: anti-Beta actin (Sigma-Aldrich, A5316), anti-Histone H3 (Cell signaling, 2650), anti-ATRX (Santa Cruz, sc-15408), anti-hMRE-11 (Calbiochem, PC388), anti-GAPDH (Cell signaling, 97166), anti-Lamin A/C (Santa Cruz, sc-376248), anti-Emerin (Santa Cruz, sc-25284), anti-RAD51AP1 (Proteintech, 11255), and anti-DAXX (Santa Cruz, sc-8043).

### RNA extraction and quantification

For RNA quantification, total RNA was extracted by using Direct-zol RNA Miniprep kits (ZYMO) according to the manufacturer’s protocol. cDNAs were synthesized using RevertAid First Strand cDNA synthesis kit (Thermo scientific) and qPCRs were performed using TB Green Premix Ex Taq II (Tli RNase H Plus) (Takara) according to the manufacturer’s protocol.

The following DNA probes were used for qPCR:

GAPDH F: TGAGAACGGGAAGCTTGTCA,

GAPDH R: AGCATCGCCCCACTTGATT,

STN1 F: TGCACAGAAAGATCCACCGG,

STN1 R: AGGCCAAGATGTGCAGGAA,

DNA2 F: ACTGTGAAAAGCGCCTGGT,

DNA2 R: AGATGTGCAGTCTCCCTCCA,

FEN1 F: TTCTTCAAGGTGACCGGCTC,

FEN1 R: TTAAACTTCCCTGCTGCCCC,

FANCM F: AGACCTTTATTGCCGCCGT,

FANCM R: TTCGTTGGGGCCATGAAGA

### Statistical Analyses

Statistical analyses for the experiments were conducted using GraphPad Prism 8. The graph displays all individual values. Detailed information regarding the tests and corresponding p-values is provided in the figures and their respective legends.

## Data Availability

For additional information and requests regarding resources or reagents, please direct your inquiries to the corresponding author, Dr. Jaewon Min. Dr. Min is responsible for handling and fulfilling these requests.

## Acknowledgements and funding sources

We thank members of the Telomere biology Lab (Min Lab) at Columbia University for their invaluable critiques, discussions, and protocols. We thank Drs. Eros Lazzerini Denchi (NIH/NCI) and Alessandro Vindigni (Washington University) for providing U2OS BLM KO and U2OS PRIMPOL KO cells. We also thank Drs. Robert Lu (CMRI), Greg Ira (Baylor College of Medicine), Kurt Runge (Cleveland Clinic), Ji-Yeon Shin (Columbia), Jean Gautier (Columbia), Max E. Gottesman (Columbia) and Shan Zha (Columbia) for key reagents and helpful discussion. This work was supported by the following NIH/NCI grants: CA207209 (RJO), CA262316 (RJO), CA197774 (AC), and CA245259 (JM).

## Notes

### Competing Interest Statement

The authors have declared no competing interest.

### Summary of Updates

Summary of Major Revisions: We have meticulously addressed all comments from the reviewers, notably by adding 42 new panels to enhance our representing evidence. We clarified that C-rich ssDNAs are nascently generated and can exist in both linear and circular forms, including both C-circles and potential precursor linear C-rich ssDNAs. We have made substantial modifications to our figures for clarity and the incorporation of pivotal new data. We determined the exact size of C-rich ssDNAs using a nucleotide ladder, revealing sizes between 200-1500nt. Additionally, Cesium density gradient analysis confirmed their ssDNA nature, and through various nucleases analyses, we gained more insights into their properties. The native IdU pulldown further showed their nascent generation. These comprehensive updates provide profound mechanistic insights, bolstering the evidence that supports our claims and significantly elevating the rigor of our initial findings. We are listing below all additional figure panels that have been added to the main figures and supplementary figures. We have also reorganized the figures. Figure 1C has been moved to Figure S1D with added western blot. Figure 1D has been added. Figure 2 (new) has 6 new panels: 2A, B, C, D, E, F Figure 3 (Previous Figure 2): 3 new panels have been added: 3E, F, G Figure 4 (Previous Figure 3): 4 panels have been revised or added: 4C, D, E, F Figure 6 (Previous Figure 5): 1 panel has been added: 6E. Figure 7 (Previous Figure 6): 2 panel have been added: 7A, B Figure S1: 12 new panels have been added: S1A, B, C, F, G, H, I, J, K, M, N, O Figure S2 (new) has 3 new panels: S2A, B, C Figure S3 (Previous Figure S2): 6 new panels have been added: S3D, E, F, G, H, I Figure S4 (Previous Figure S3): 2 panels have been revised: S4B, E Figure S5 (Previous Figure S4): 5 new panels have been added: S5C, D, F Figure S6 (Previous Figure S5): 2 new panels have been added: S6F, G

## References

1. J. W. Shay, W. E. Wright, Telomeres and telomerase: three decades of progress. Nat Rev Genet 20, 299–309 (2019).

2. T. M. Bryan, A. Englezou, L. Dalla-Pozza, M. A. Dunham, R. R. Reddel, Evidence for an alternative mechanism for maintaining telomere length in human tumors and tumor-derived cell lines. Nat Med 3, 1271–1274 (1997).

3. J. R. Lydeard, S. Jain, M. Yamaguchi, J. E. Haber, Break-induced replication and telomerase-independent telomere maintenance require Pol32. Nature 448, 820–823 (2007).

4. F. M. Roumelioti et al., Alternative lengthening of human telomeres is a conservative DNA replication process with features of break-induced replication. EMBO Rep 17, 1731–1737 (2016).

5. R. L. Dilley et al., Break-induced telomere synthesis underlies alternative telomere maintenance. Nature 539, 54–58 (2016).

6. J. Min, W. E. Wright, J. W. Shay, Alternative Lengthening of Telomeres Mediated by Mitotic DNA Synthesis Engages Break-Induced Replication Processes. Mol Cell Biol 37 (2017).

7. A. J. Cesare, R. R. Reddel, Alternative lengthening of telomeres: models, mechanisms and implications. Nat Rev Genet 11, 319–330 (2010).

8. A. J. Cesare, J. D. Griffith, Telomeric DNA in ALT cells is characterized by free telomeric circles and heterogeneous t-loops. Mol Cell Biol 24, 9948–9957 (2004).

9. Y. Doksani, J. Y. Wu, T. de Lange, X. Zhuang, Super-resolution fluorescence imaging of telomeres reveals TRF2-dependent T-loop formation. Cell 155, 345–356 (2013).

10. G. Mazzucco et al., Telomere damage induces internal loops that generate telomeric circles. Nat Commun 11, 5297(2020).

11. T. Zhang et al., Looping-out mechanism for resolution of replicative stress at telomeres. EMBO Rep 18, 1412–1428 (2017).

12. H. A. Pickett, A. J. Cesare, R. L. Johnston, A. A. Neumann, R. R. Reddel, Control of telomere length by a trimming mechanism that involves generation of t-circles. EMBO J 28, 799–809 (2009).

13. J. S. Li et al., TZAP: A telomere-associated protein involved in telomere length control. Science 355, 638–641 (2017).

14. H. A. Pickett, J. D. Henson, A. Y. Au, A. A. Neumann, R. R. Reddel, Normal mammalian cells negatively regulate telomere length by telomere trimming. Hum Mol Genet 20, 4684–4692 (2011).

15. L. Oganesian, J. Karlseder, 5’ C-rich telomeric overhangs are an outcome of rapid telomere truncation events. DNA Repair (Amst) 12, 238–245 (2013).

16. J. D. Henson et al., DNA C-circles are specific and quantifiable markers of alternative-lengthening-of-telomeres activity. Nat Biotechnol 27, 1181–1185 (2009).

17. A. I. Idilli, S. Segura-Bayona, T. P. Lippert, S. J. Boulton, A C-circle assay for detection of alternative lengthening of telomere activity in FFPE tissue. STAR Protoc 2, 100569 (2021).

18. Y. Y. Chen et al., The C-Circle Biomarker Is Secreted by Alternative-Lengthening-of-Telomeres Positive Cancer Cells inside Exosomes and Provides a Blood-Based Diagnostic for ALT Activity. Cancers (Basel) 13 (2021).

19. J. D. Henson et al., The C-Circle Assay for alternative-lengthening-of-telomeres activity. Methods 114, 74–84 (2017).

20. C. Y. Jones et al., Hyperextended telomeres promote formation of C-circle DNA in telomerase positive human cells. J Biol Chem 299, 104665 (2023).

21. J. Min, W. E. Wright, J. W. Shay, Alternative lengthening of telomeres can be maintained by preferential elongation of lagging strands. Nucleic Acids Res 45, 2615–2628 (2017).

22. T. Zhang et al., Strand break-induced replication fork collapse leads to C-circles, C-overhangs and telomeric recombination. PLoS Genet 15, e1007925 (2019).

23. R. Lu, H. A. Pickett, Telomeric replication stress: the beginning and the end for alternative lengthening of telomeres cancers. Open Biol 12, 220011 (2022).

24. A. Nabetani, F. Ishikawa, Unusual telomeric DNAs in human telomerase-negative immortalized cells. Mol Cell Biol 29, 703–713 (2009).

25. L. Oganesian, J. Karlseder, Mammalian 5’ C-rich telomeric overhangs are a mark of recombination-dependent telomere maintenance. Mol Cell 42, 224–236 (2011).

26. T. Zhang et al., Break-induced replication orchestrates resection-dependent template switching. Nature 619, 201–208 (2023).

27. Y. K. Kim, J. Yeo, B. Kim, M. Ha, V. N. Kim, Short structured RNAs with low GC content are selectively lost during extraction from a small number of cells. Mol Cell 46, 893–895 (2012).

28. T. P. Lai, W. E. Wright, J. W. Shay, Generation of digoxigenin-incorporated probes to enhance DNA detection sensitivity. Biotechniques 60, 306–309 (2016).

29. G. D. Birnie, Separation of native and denatured DNA, RNA and hybrid on sodium iodide gradients. FEBS Lett 27, 19–22 (1972).

30. A. Dupre et al., A forward chemical genetic screen reveals an inhibitor of the Mre11-Rad50-Nbs1 complex. Nat Chem Biol 4, 119–125 (2008).

31. R. J. O’Sullivan et al., Rapid induction of alternative lengthening of telomeres by depletion of the histone chaperone ASF1. Nat Struct Mol Biol 21, 167–174 (2014).

32. A. Shibata et al., DNA double-strand break repair pathway choice is directed by distinct MRE11 nuclease activities. Mol Cell 53, 7–18 (2014).

33. N. Arnoult, C. Saintome, I. Ourliac-Garnier, J. F. Riou, A. Londono-Vallejo, Human POT1 is required for efficient telomere C-rich strand replication in the absence of WRN. Genes Dev 23, 2915–2924 (2009).

34. J. Zimmer et al., Targeting BRCA1 and BRCA2 Deficiencies with G-Quadruplex-Interacting Compounds. Mol Cell 61, 449–460 (2016).

35. M. Zimmermann, T. Kibe, S. Kabir, T. de Lange, TRF1 negotiates TTAGGG repeat-associated replication problems by recruiting the BLM helicase and the TPP1/POT1 repressor of ATR signaling. Genes Dev 28, 2477–2491 (2014).

36. J. Lee et al., Dynamic interaction of BRCA2 with telomeric G-quadruplexes underlies telomere replication homeostasis. Nat Commun 13, 3396 (2022).

37. W. Lin et al., Mammalian DNA2 helicase/nuclease cleaves G-quadruplex DNA and is required for telomere integrity. EMBO J 32, 1425–1439 (2013).

38. H. Sun et al., Okazaki fragment maturation: DNA flap dynamics for cell proliferation and survival. Trends Cell Biol 33, 221–234 (2023).

39. H. Hanzlikova et al., The Importance of Poly(ADP-Ribose) Polymerase as a Sensor of Unligated Okazaki Fragments during DNA Replication. Mol Cell 71, 319–331 e313 (2018).

40. K. Cong et al., Replication gaps are a key determinant of PARP inhibitor synthetic lethality with BRCA deficiency. Mol Cell 81, 3128–3144 e3127 (2021).

41. Y. Shen et al., BMN 673, a novel and highly potent PARP1/2 inhibitor for the treatment of human cancers with DNA repair deficiency. Clin Cancer Res 19, 5003–5015 (2013).

42. S. Garcia-Gomez et al., PrimPol, an archaic primase/polymerase operating in human cells. Mol Cell 52, 541–553 (2013).

43. A. Taglialatela et al., REV1-Polzeta maintains the viability of homologous recombination-deficient cancer cells through mutagenic repair of PRIMPOL-dependent ssDNA gaps. Mol Cell 81, 4008–4025 e4007 (2021).

44. S. Tirman et al., Temporally distinct post-replicative repair mechanisms fill PRIMPOL-dependent ssDNA gaps in human cells. Mol Cell 81, 4026–4040 e4028 (2021).

45. A. Quinet et al., PRIMPOL-Mediated Adaptive Response Suppresses Replication Fork Reversal in BRCA-Deficient Cells. Mol Cell 77, 461–474 e469 (2020).

46. K. P. M. Mehta et al., CHK1 phosphorylates PRIMPOL to promote replication stress tolerance. Sci Adv 8, eabm0314 (2022).

47. C. A. Lovejoy et al., Loss of ATRX, genome instability, and an altered DNA damage response are hallmarks of the alternative lengthening of telomeres pathway. PLoS Genet 8, e1002772 (2012).

48. C. M. Heaphy et al., Altered telomeres in tumors with ATRX and DAXX mutations. Science 333, 425 (2011).

49. D. Clynes et al., Suppression of the alternative lengthening of telomere pathway by the chromatin remodelling factor ATRX. Nat Commun 6, 7538 (2015).

50. D. T. Nguyen et al., The chromatin remodelling factor ATRX suppresses R-loops in transcribed telomeric repeats. EMBO Rep 18, 914–928 (2017).

51. Y. C. Teng et al., ATRX promotes heterochromatin formation to protect cells from G-quadruplex DNA-mediated stress. Nat Commun 12, 3887 (2021).

52. X. Pan et al., FANCM suppresses DNA replication stress at ALT telomeres by disrupting TERRA R-loops. Sci Rep 9, 19110 (2019).

53. B. Silva et al., FANCM limits ALT activity by restricting telomeric replication stress induced by deregulated BLM and R-loops. Nat Commun 10, 2253 (2019).

54. R. Lu et al., The FANCM-BLM-TOP3A-RMI complex suppresses alternative lengthening of telomeres (ALT). Nat Commun 10, 2252 (2019).

55. J. Barroso-Gonzalez et al., RAD51AP1 Is an Essential Mediator of Alternative Lengthening of Telomeres. Mol Cell 76, 11–26 e17 (2019).

56. A. Decottignies, TERRA and RAD51AP1 at the R&D-loop department of ALT telomeres. Mol Cell 82, 3963–3965 (2022).

57. T. Yadav et al., TERRA and RAD51AP1 promote alternative lengthening of telomeres through an R- to D-loop switch. Mol Cell 82, 3985–4000 e3984 (2022).

58. N. Kaminski et al., RAD51AP1 regulates ALT-HDR through chromatin-directed homeostasis of TERRA. Mol Cell 82, 4001–4017 e4007 (2022).

59. C. A. Lovejoy, K. Takai, M. S. Huh, D. J. Picketts, T. de Lange, ATRX affects the repair of telomeric DSBs by promoting cohesion and a DAXX-dependent activity. PLoS Biol 18, e3000594 (2020).

60. H. E. B. Geiller et al., ATRX modulates the escape from a telomere crisis. PLoS Genet 18, e1010485 (2022).

61. C. E. Napier et al., ATRX represses alternative lengthening of telomeres. Oncotarget 6, 16543–16558 (2015).

62. S. W. Cai, T. de Lange, CST-Polalpha/Primase: the second telomere maintenance machine. Genes Dev 10.1101/gad.350479.123 (2023).

63. M. Zhang et al., Mammalian CST averts replication failure by preventing G-quadruplex accumulation. Nucleic Acids Res 47, 5243–5259 (2019).

64. Y. Miyake et al., RPA-like mammalian Ctc1-Stn1-Ten1 complex binds to single-stranded DNA and protects telomeres independently of the Pot1 pathway. Mol Cell 36, 193–206 (2009).

65. F. Wang et al., Human CST has independent functions during telomere duplex replication and C-strand fill-in. Cell Rep 2, 1096–1103 (2012).

66. C. Huang, P. Jia, M. Chastain, O. Shiva, W. Chai, The human CTC1/STN1/TEN1 complex regulates telomere maintenance in ALT cancer cells. Exp Cell Res 355, 95–104 (2017).

67. C. Huang, X. Dai, W. Chai, Human Stn1 protects telomere integrity by promoting efficient lagging-strand synthesis at telomeres and mediating C-strand fill-in. Cell Res 22, 1681–1695 (2012).

68. J. Min, W. E. Wright, J. W. Shay, Clustered telomeres in phase-separated nuclear condensates engage mitotic DNA synthesis through BLM and RAD52. Genes Dev 33, 814–827 (2019).

69. A. P. Sobinoff et al., BLM and SLX4 play opposing roles in recombination-dependent replication at human telomeres. EMBO J 36, 2907–2919 (2017).

70. T. K. Loe et al., Telomere length heterogeneity in ALT cells is maintained by PML-dependent localization of the BTR complex to telomeres. Genes Dev 34, 650–662 (2020).

71. S. Li et al., PIF1 helicase promotes break-induced replication in mammalian cells. EMBO J 40, e104509 (2021).

72. J. Min et al., Mechanisms of insertions at a DNA double-strand break. Mol Cell 10.1016/j.molcel.2023.06.016 (2023).

73. L. S. Symington, J. Gautier, Double-strand break end resection and repair pathway choice. Annu Rev Genet 45, 247–271 (2011).

74. J. Oh, L. S. Symington, Role of the Mre11 Complex in Preserving Genome Integrity. Genes (Basel) 9 (2018).

75. V. Costanzo et al., Mre11 protein complex prevents double-strand break accumulation during chromosomal DNA replication. Mol Cell 8, 137–147 (2001).

76. B. M. Sirbu et al., Analysis of protein dynamics at active, stalled, and collapsed replication forks. Genes Dev 25, 1320–1327 (2011).

77. R. S. Maser et al., Mre11 complex and DNA replication: linkage to E2F and sites of DNA synthesis. Mol Cell Biol 21, 6006–6016 (2001).

78. R. Rai et al., NBS1 Phosphorylation Status Dictates Repair Choice of Dysfunctional Telomeres. Mol Cell 65, 801–817 e804 (2017).

79. X. D. Zhu, B. Kuster, M. Mann, J. H. Petrini, T. de Lange, Cell-cycle-regulated association of RAD50/MRE11/NBS1 with TRF2 and human telomeres. Nat Genet 25, 347–352 (2000).

80. S. A. Compton, J. H. Choi, A. J. Cesare, S. Ozgur, J. D. Griffith, Xrcc3 and Nbs1 are required for the production of extrachromosomal telomeric circles in human alternative lengthening of telomere cells. Cancer Res 67, 1513–1519 (2007).

81. Z. H. Zhong et al., Disruption of telomere maintenance by depletion of the MRE11/RAD50/NBS1 complex in cells that use alternative lengthening of telomeres. J Biol Chem 282, 29314–29322 (2007).

82. E. Y. Yu, J. Perez-Martin, W. K. Holloman, N. F. Lue, Mre11 and Blm-Dependent Formation of ALT-Like Telomeres in Ku-Deficient Ustilago maydis. PLoS Genet 11, e1005570 (2015).

83. W. Q. Jiang et al., Suppression of alternative lengthening of telomeres by Sp100-mediated sequestration of the MRE11/RAD50/NBS1 complex. Mol Cell Biol 25, 2708–2721 (2005).

84. J. W. Leung et al., Alpha thalassemia/mental retardation syndrome X-linked gene product ATRX is required for proper replication restart and cellular resistance to replication stress. J Biol Chem 288, 6342–6350 (2013).

85. S. Moreau, J. R. Ferguson, L. S. Symington, The nuclease activity of Mre11 is required for meiosis but not for mating type switching, end joining, or telomere maintenance. Mol Cell Biol 19, 556–566 (1999).

86. T. T. Paull, M. Gellert, The 3’ to 5’ exonuclease activity of Mre 11 facilitates repair of DNA double-strand breaks. Mol Cell 1, 969–979 (1998).

87. S. Panier et al., SLX4IP Antagonizes Promiscuous BLM Activity during ALT Maintenance. Mol Cell 76, 27–43 e11 (2019).

88. Y. Yu et al., Dna2 nuclease deficiency results in large and complex DNA insertions at chromosomal breaks. Nature 564, 287–290 (2018).

89. C. Zhou, S. Pourmal, N. P. Pavletich, Dna2 nuclease-helicase structure, mechanism and regulation by Rpa. Elife 4 (2015).

90. T. Li et al., Cooperative interaction of CST and RECQ4 resolves G-quadruplexes and maintains telomere stability. EMBO Rep 10.15252/embr.202255494, e55494 (2023).

91. A. Emam et al., Stalled replication fork protection limits cGAS-STING and P-body-dependent innate immune signalling. Nat Cell Biol 24, 1154–1164 (2022).

92. I.-C. Cheng et al., Wuho is a new member in maintaining genome stability through its interaction with flap endonuclease 1. PLoS biology 14, e1002349 (2016).

93. W. Chai, A. J. Sfeir, H. Hoshiyama, J. W. Shay, W. E. Wright, The involvement of the Mre11/Rad50/Nbs1 complex in the generation of G-overhangs at human telomeres. EMBO Rep 7, 225–230 (2006).

