## Supporting information S1_S6 for "Extrachromosomal Telomeres Derived from Excessive Strand Displacements"

**Figure S1, related to Figure 1. 4SET assay, a simple and efficient method for detecting single-stranded extrachromosomal telomeric DNA. (A)** Comparison of genomic DNA purification between DNeasy Blood & Tissue Kit (Qiagen, 69504) and our 4SET method in U2OS and Saos2 cells. **(B)** Comparison between Dig-labeled strand-specific probes and standard end-labeling probes. **(C)** Comparison of 4SET results with DNA dissolved at 4°C overnight vs 55°C for 2 hours, followed by 4°C overnight. Observed smear signals potentially indicate the presence of nicked or gapped DNA. The graph to the right displays the quantified signals potentially indicative of nicked or gapped DNA (mean ± SD; unpaired t-test). **(D)** Western blot analysis of the whole, chromatin (Chr.), and woluble using anti-Histone H3, Lamin A/C, Emerin, GAPDH, and beta-actin for the validation of cell fractions. **(E)** SYBR DNA images for G-rich probe and C-rich probe gels in Fig. 1C. **(F)** Western blot analysis of the whole, chromatin (Chr.), and soluble fractions using PML antibody. **(G)** Western blot analysis of Saos2 Wild-type and PML Knock-out cells using PML antibody, along with a Ponceau image as a loading control. **(H)** 4SET assay for SaoS2 Wild-type and PML Knock-out cells after fractionation into Whole and soluble fractions. G-rich probe was used to detect C-rich telomeric sequences, and C-rich probe was used to detect G-rich telomeric sequences. C-rich single stranded DNAs are indicated by an arrow. SYBR DNA staining of gel image. **(I)** C-circle assay using DNA in (H) (top). Quantification of C-circle assay (mean ± SD; unpaired t-test) (bottom). **(J)** 4SET assay conducted using a nucleotide ladder (RiboRuler) to measure the size of C-rich single-stranded DNA in SaoS2 soluble fraction, analyzed on both 0.5% and 1% agarose gels. **(K)** 4SET assay for SaoS2 treated with Zeocin (100 μg/ml), Etoposide (ETO) (10 μM), or Camptothecin (CPT) (0.25 μM) for 24 hrs. **(L)** Quantification of the C-circle assay in Fig. 1E (SaoS2 cells). Relative amount of C-circle assay products (mean ± SD; unpaired t-test). **(M)** Cell fractionation using E1 and E2 buffers to separate chromatin, nucleoplasm, and cytoplasm. **(N)** Western blot analysis conducted using the fractionated DNAs in (M) using anti-Histone H3, Lamin A/C, GAPDH, and PML for the validation of cell fractions. **(O)** 4SET assay for fractionated DNAs in (M).

**Figure S2, related to Figure 2. The presence of C-rich telomeric single-stranded DNA. (A)** S1 nuclease assay on U2OS and SaoS2 soluble DNA fractions. **(B)** Lambda exonuclease assay on U2OS soluble DNA fraction. **(C)** Telomere-FISH with TelG (G-rich) and TelC (C-rich) PNA probes for chromatin (denatured FISH) and soluble fractions (native FISH).

**Figure S3, related to Figure 3. MRE11 nuclease activity suppresses the generation of C-rich ssDNA. (A)** SYBR DNA image for G-rich probe and C-rich probe gels in Fig. 3D. **(B)** 4SET assay for HeLa LT cells with or without Mirin treatment. Samples were fractionated into soluble DNA fraction or whole DNA. G-rich probe was used to detect C-rich telomeric sequences, and C-rich probe was used to detect G-rich telomeric sequences. SYBR DNA image for G-probe gel. **(C)** 4SET assay for SaoS2 cells with Mirin, PFM01, PFM39 or mock (control) treatments. G-rich probe was used to detect C-rich telomeric sequences, and C-rich probe was used to detect G-rich telomeric sequences. C-rich single stranded DNAs are indicated by an arrow. SYBR DNA image for G-rich probe and C-rich probe gels. **(D)** Western blot of SaoS2 post-transfection with siLuciferase (siLuc) or siMRE11s. **(E)** 4SET for SaoS2, with and without Mre11 treatment post siLuc or siMRE11s transfection. The graph to the right presents the quantification of relative amount of C-rich ssDNAs (mean ± SD; unpaired t-test). **(F)** 4SET for SaoS2 WT and SaoS2 PML KO with/without Mirin treatment. **(G)** Quantification of 4SET from F, showing relative C-rich ssDNA (whole) levels (mean ± SD). **(H)** Native IdU pulldown was conducted using IdU (3D4) antibody for Saos2 WT and Saos2 PML KO after IdU incorporation. IgG served as a negative control. Chromatin DNA was also included to control for DNA quantity. **(I)** Quantification of the native IdU-pulldown assay in H; Relative amount of nascent C-rich ssDNAs (mean ± SD; unpaired t-test).

**Figure S4, related to Figure 4. C-rich ssDNAs are derived from lagging strand during the Okazaki fragment processing.** **(A)** Relative amount of DNA2 mRNA levels measured by quantitative-PCR in U2OS cells transfected with siRNAs targeting DNA2 or control (siLuc). **(B)** SYBR DNA images for G-rich probe and C-rich probe gels in Fig. 4C. **(C)** Relative amount of FEN1 mRNA levels measured by quantitative-PCR in U2OS cells transfected with siRNAs targeting FEN1 or control (siLuc). **(D)** Illustration depicting 5' flap processing at lagging strand telomeres by DNA2-RPA or FEN1-PCNA. **(E)** SYBR DNA images for G-rich probe and C-rich probe gels in Fig. 4E. **(F)** 4SET assay for U2OS cells with PARPi (Talazoparib), Mirin, or mock (control) treatments. G-rich probe was used to detect C-rich telomeric sequences, and C-rich probe was used to detect G-rich telomeric sequences. C-rich single stranded DNAs are indicated by an arrow. SYBR DNA images for G-rich probe and C-rich probe gels. **(G)** 4SET assay for SaoS2 cells after transfection of siRNAs targeting PRIMPOL, or control (siLuc) with or without Mirin treatment. G-rich probe was used to detect C-rich telomeric sequences. C-rich single stranded DNAs are indicated by an arrow. SYBR DNA image for G-rich probe. **(H)** 4SET assay for U2OS PRIMPOL KO cells after transfection of PRIMPOL WT, S255D, or control cDNA plasmid with or without Mirin treatment. G-rich probe was used to detect C-rich telomeric sequences, and C-rich probe was used to detect G-rich telomeric sequences. C-rich single stranded DNAs are indicated by an arrow. SYBR DNA images for G-rich probe and C-rich probe gels.

**Figure S5, related to Figure 5. ATRX suppresses the generation of C-rich ssDNAs.** **(A)** SYBR DNA images for G-rich probe and C-rich probe gels in Fig. 5D. **(B)** SYBR DNA images for G-rich probe and C-rich probe gels in Fig. 5G. **(C)** Relative amount of FANCM mRNA levels measured by quantitative-PCR in U2OS cells transfected with siRNAs targeting FANCM or control (siLuc). **(D)** Relative quantification of R-loops at telomeres using the 7p TERRA primer following DNA/RNA hybrid immunoprecipitation with the S9.6 antibody or IgG in U2OS cells transfected with siRNAs targeting FANCM or control (siLuc). **(E)** 4SET assay for U2OS cells after transfection of siRNAs targeting FANCM or control (siLuc) with Mirin treatment. G-rich probe was used to detect C-rich telomeric sequences. C-rich single stranded DNAs are indicated by an arrow. SYBR DNA image for G-rich probe gel. **(F)** Western blot analysis of U2OS RAD51AP1 KO (C1) or control. **(G)** 4SET assay for U2OS RAD51AP1 KO (C1) or control with Mirin treatment. G-rich probe was used to detect C-rich telomeric sequences, and C-rich probe was used to detect G-rich telomeric sequences. C-rich single stranded DNAs are indicated by an arrow. SYBR DNA image for G-rich probe gel. **(H)** 4SET assay for HeLa LT TERC KO or HeLa LT TERC KO cells expressing shRNA targeting ATRX gene (ATRX 590) with or without Mirin or PARPi treatment. G-rich probe was used to detect C-rich telomeric sequences, and C-rich probe was used to detect G-rich telomeric sequences. SYBR DNA image for G-rich probe gel.

**Figure S6, related to Figure 6. CST complex-mediated priming and subsequent strand displacements by DNA polymerase delta generates C-rich ssDNAs. (A)** Relative amount of STN1 mRNA levels measured by quantitative-PCR in U2OS and SaoS2 cells expressing shRNA targeting STN1 gene or control (shNT). **(B)** SYBR DNA images for G-rich probe and C-rich probe gels in Fig. 6D. **(C)** 4SET assay for U2OS cells expressing shRNAs targeting STN1, or non-targeting (NT) control. Cells were treated with Mirin (50 μM, 48 hr). SYBR DNA images for G-rich probe and C-rich probe gels. **(D)** SYBR DNA images for G-rich probe and C-rich probe gels in Fig. 6F. **(E)** 4SET assay for Mirin treated U2OS cells along with aphidicolin, CD437 or control. Samples were fractionated into soluble DNA fraction or whole DNA. G-rich probe was used to detect C-rich telomeric sequences, and C-rich probe was used to detect G-rich telomeric sequences. C-rich single stranded DNAs are indicated by an arrow. SYBR DNA image for G-rich probe gel. **(F)** 4SET assay for Mirin treated U2OS WT or BLM KO cells. G-rich probe was used to detect C-rich telomeric sequences, and C-rich probe was used to detect G-rich telomeric sequences. C-rich single stranded DNAs are indicated by an arrow. SYBR DNA image for G-rich probe gel. **(G)** Quantification of the 4SET assay in F; as the relative amount of C-rich ssDNA (whole). (mean ± SD; unpaired t-test). **(H)** SYBR DNA images for G-rich probe and C-rich probe gels in Fig. 6G.

### Figure S1

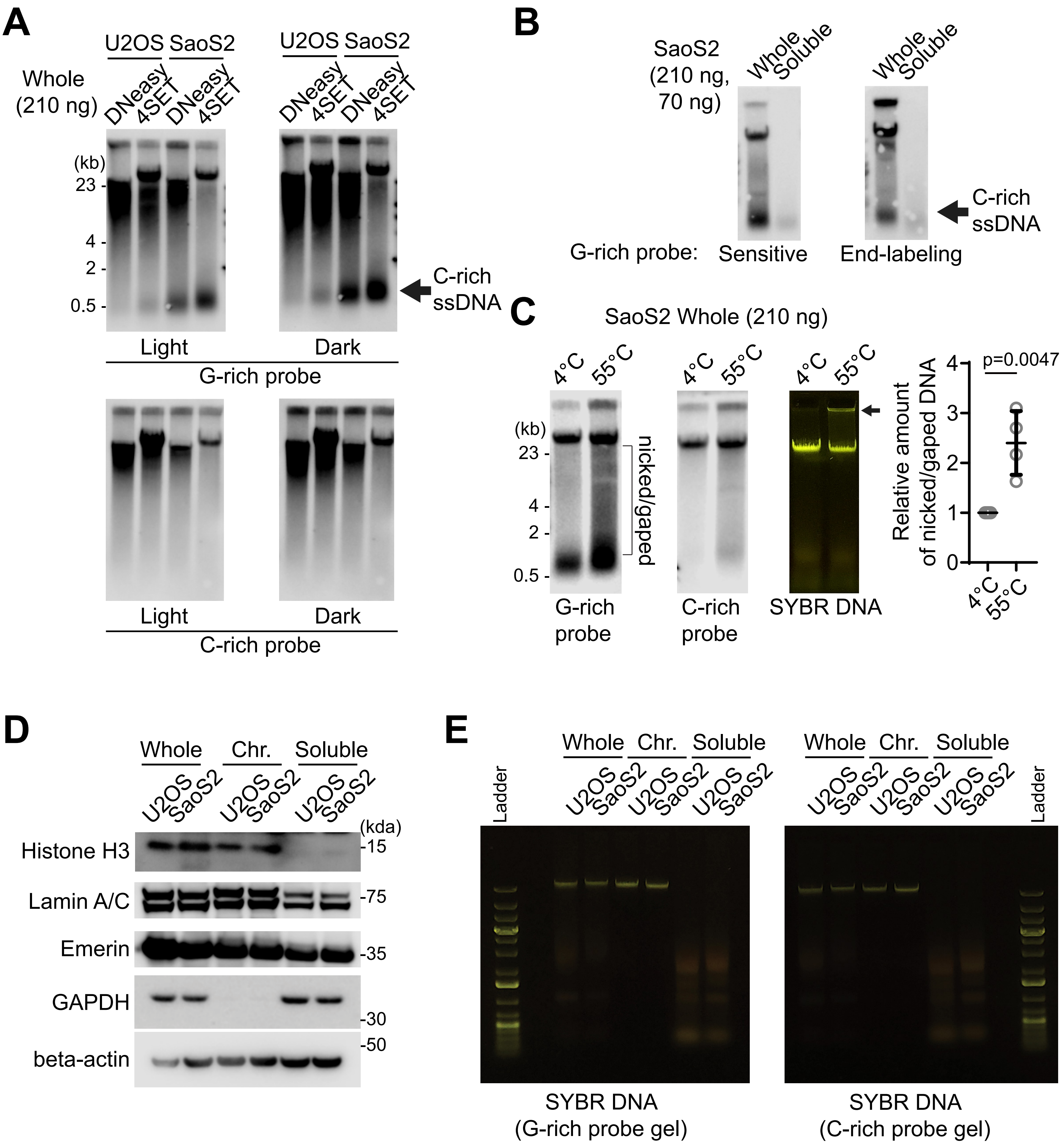
A
U2OS SaoS2 U2OS SaoS2
Whole (210 ng)
DNeasy 4SET DNeasy 4SET DNeasy 4SET DNeasy 4SET
(kb)
23
4
2
0.5
C-rich ssDNA
Light
Dark
G-rich probe
Light
Dark
C-rich probe
B
SaoS2 (210 ng, 70 ng)
Whole Soluble
Whole Soluble
C-rich ssDNA
G-rich probe:
Sensitive
End-labeling
C
SaoS2 Whole (210 ng)
4°C 55°C
(kb)
23
4
2
0.5
nicked/gaped
G-rich probe
C-rich probe
SYBR DNA
p=0.0047
Relative amount of nicked/gaped DNA
0
1
2
3
4
D
Whole Chr. Soluble
U2OS SaoS2 U2OS SaoS2 U2OS SaoS2
(kda)
Histone H3
-15
Lamin A/C
-75
Emerin
-35
GAPDH
-30
-50
beta-actin
E
Whole Chr. Soluble
U2OS SaoS2 U2OS SaoS2 U2OS SaoS2
Ladder
SYBR DNA (G-rich probe gel)
SYBR DNA (C-rich probe gel)

### Figure S1

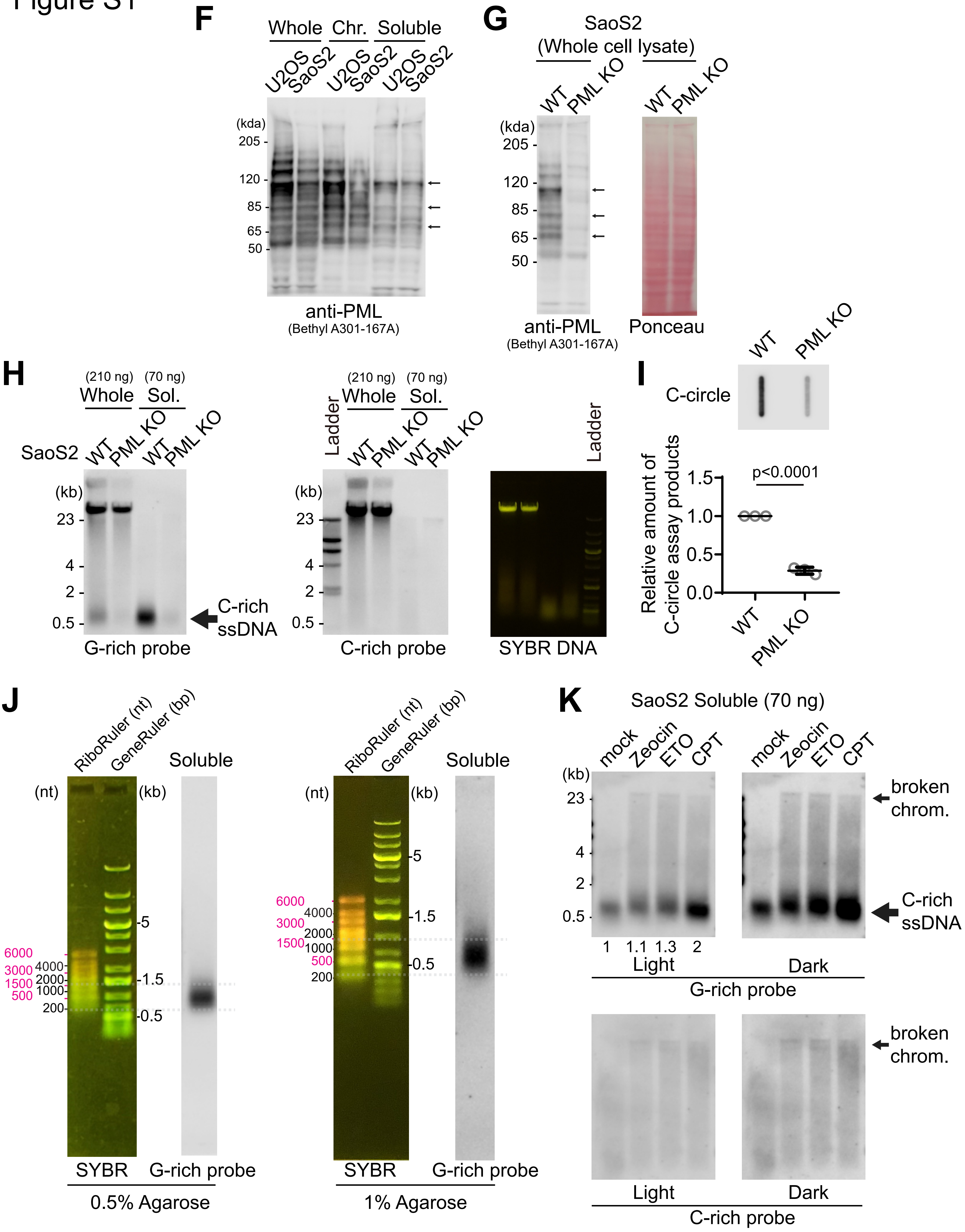
F
Whole
Chr.
Soluble
U2OS
SaoS2
U2OS
SaoS2
U2OS
SaoS2
(kda)
205
120
85
65
50
anti-PML
(Bethyl A301-167A)
G
SaoS2
(Whole cell lysate)
WT
PML KO
WT
PML KO
(kda)
205
120
85
65
50
anti-PML
(Bethyl A301-167A)
Ponceau
H
(210 ng)
Whole
(70 ng)
Sol.
SaoS2
WT
PML KO
WT
PML KO
(kb)
23
4
2
0.5
C-rich
ssDNA
G-rich probe
Ladder
(210 ng)
Whole
(70 ng)
Sol.
WT
PML KO
WT
PML KO
(kb)
23
4
2
0.5
C-rich probe
Ladder
SYBR DNA
I
WT
PML KO
C-circle
Relative amount of
C-circle assay products
p<0.0001
1.5
1.0
0.5
0.0
WT
PML KO
J
RiboRuler (nt)
GeneRuler (bp)
Soluble
(nt)
(kb)
6000
3000
1500
500
4000
2000
1000
200
5
1.5
0.5
SYBR
G-rich probe
0.5% Agarose
RiboRuler (nt)
GeneRuler (bp)
Soluble
(nt)
(kb)
6000
3000
1500
500
4000
2000
1000
200
5
1.5
0.5
SYBR
G-rich probe
1% Agarose
K
SaoS2 Soluble (70 ng)
mock
Zeocin
ETO
CPT
mock
Zeocin
ETO
CPT
(kb)
23
4
2
0.5
broken
chrom.
C-rich
ssDNA
1
1.1
1.3
2
Light
Dark
G-rich probe
broken
chrom.
Light
Dark
C-rich probe

### Figure S1

**L**

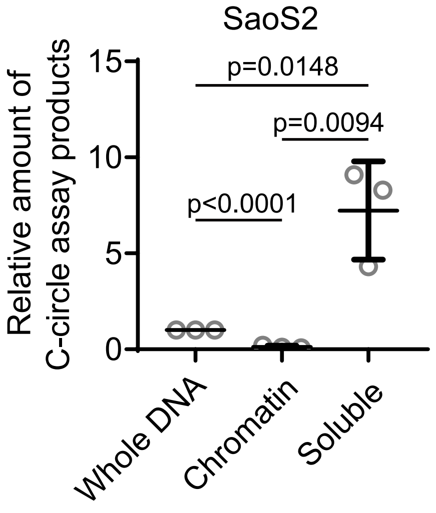


**M**

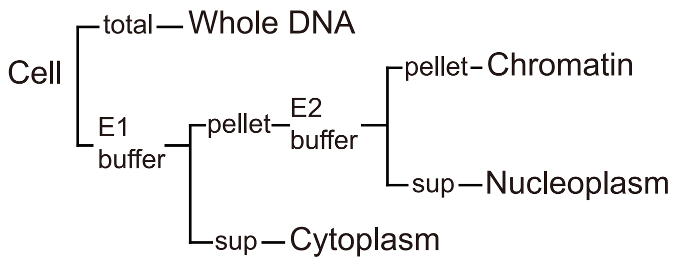


**N**

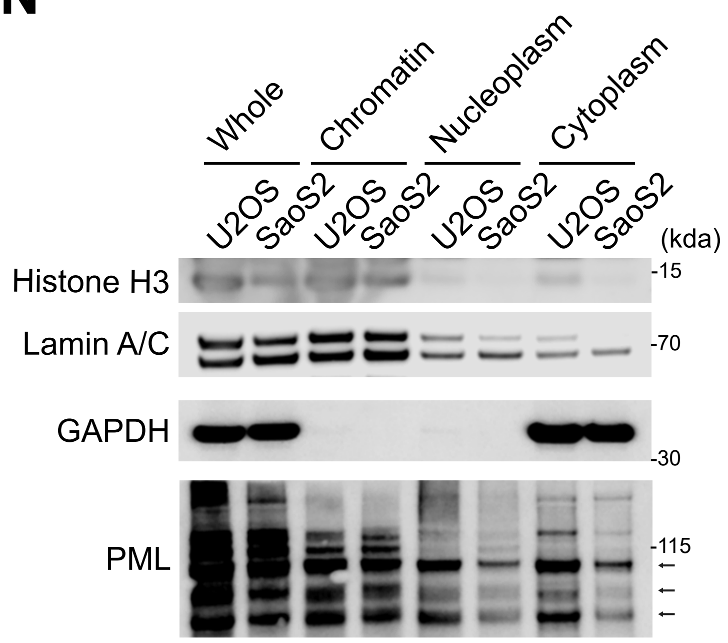


**O**

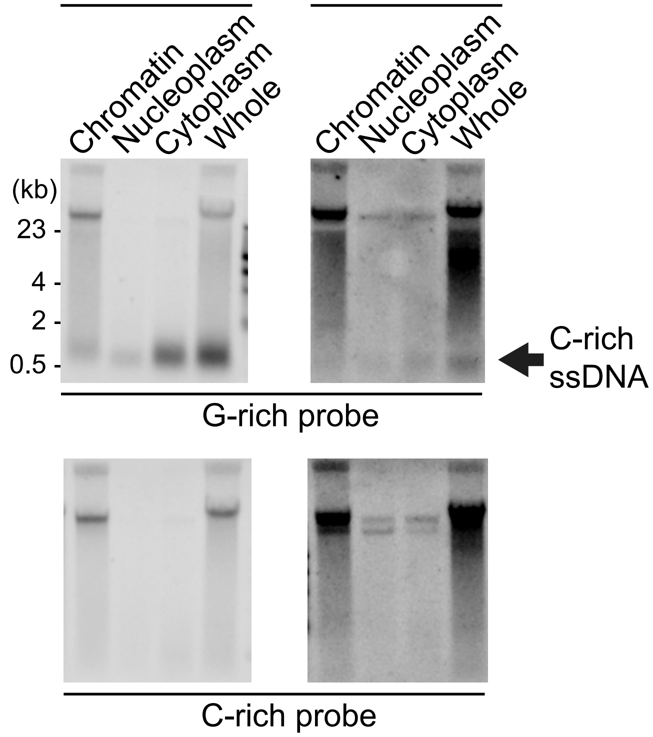


Whole - 140ng
Chromatin&Nucleoplasm&Cytoplasm - 70ng

### Figure S2

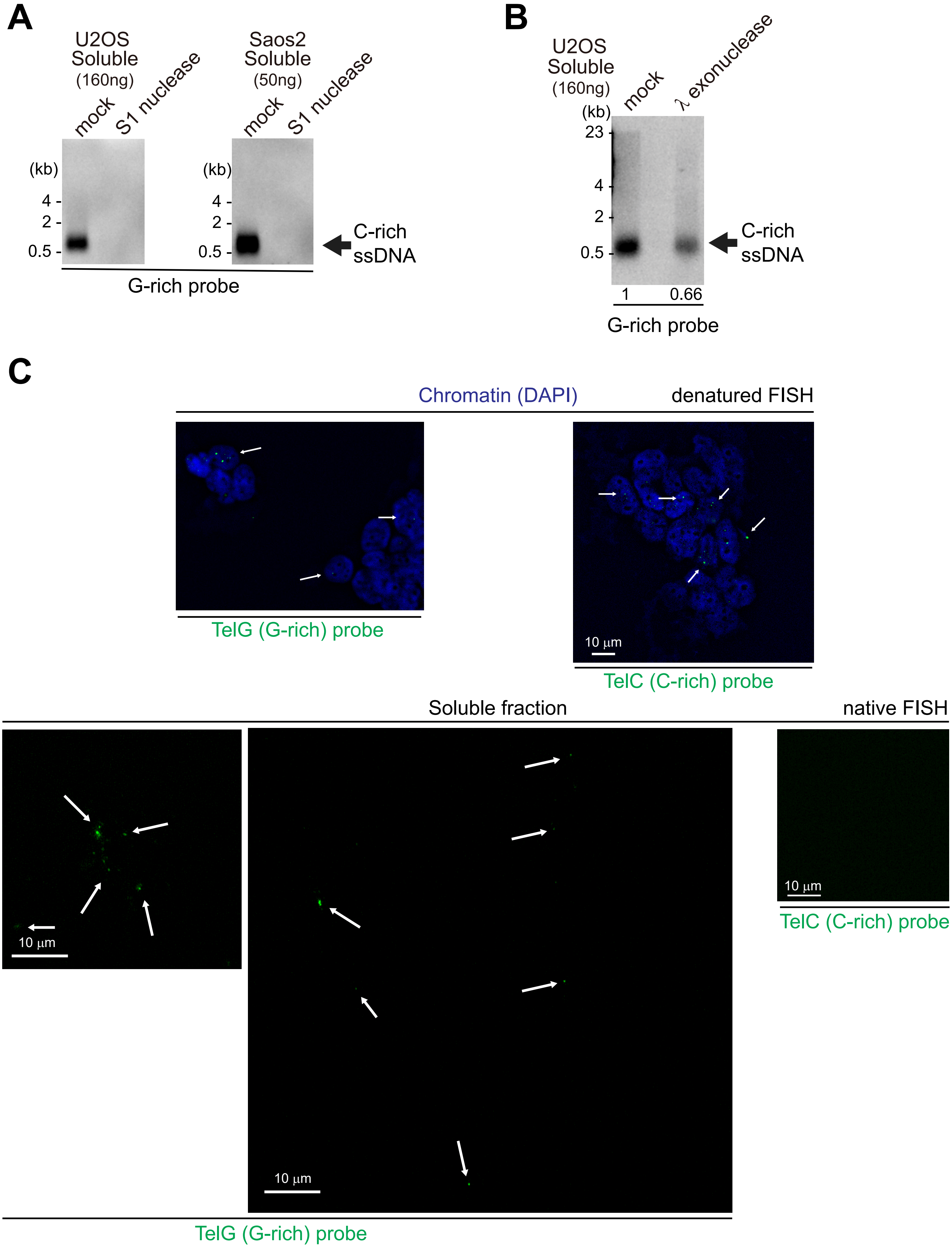

A
U2OS
Soluble
(160ng)
mock
S1 nuclease
Saos2
Soluble
(50ng)
mock
S1 nuclease
(kb)
4
2
0.5
C-rich
ssDNA
G-rich probe
B
U2OS
Soluble
(160ng)
mock
λ exonuclease
(kb)
23
4
2
0.5
C-rich
ssDNA
1
0.66
G-rich probe
C
Chromatin (DAPI)
denatured FISH
10 μm
TelG (G-rich) probe
TelC (C-rich) probe
Soluble fraction
native FISH
10 μm
10 μm
10 μm
TelC (C-rich) probe
TelG (G-rich) probe

### Figure S3

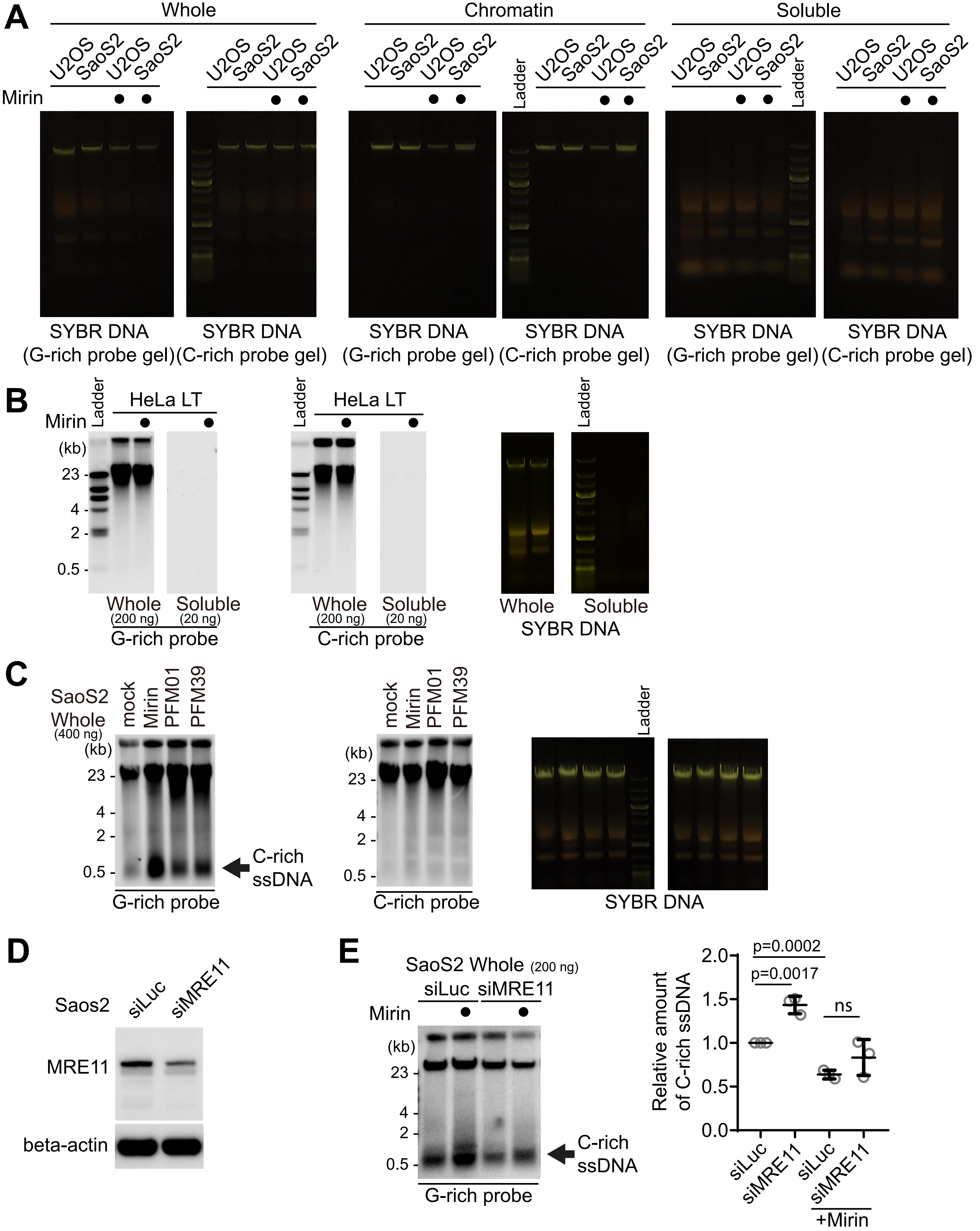
A
Whole
Chromatin
Soluble
U2OS SaoS2 U2OS SaoS2
Mirin
Ladder
SYBR DNA (G-rich probe gel)
SYBR DNA (C-rich probe gel)
B
HeLa LT
Ladder
(kb)
23
4
2
0.5
Whole (200 ng)
Soluble (20 ng)
G-rich probe
C-rich probe
Whole
Soluble
SYBR DNA
C
SaoS2 Whole (400 ng)
mock Mirin PFM01 PFM39
C-rich ssDNA
G-rich probe
C-rich probe
SYBR DNA
D
Saos2
siLuc siMRE11
MRE11
beta-actin
E
SaoS2 Whole (200 ng)
siLuc siMRE11
Mirin
C-rich ssDNA
G-rich probe
p=0.0002
p=0.0017
ns
Relative amount of C-rich ssDNA
2.0
1.5
1.0
0.5
0.0
siLuc siMRE11 siLuc siMRE11
+Mirin

Figure S3

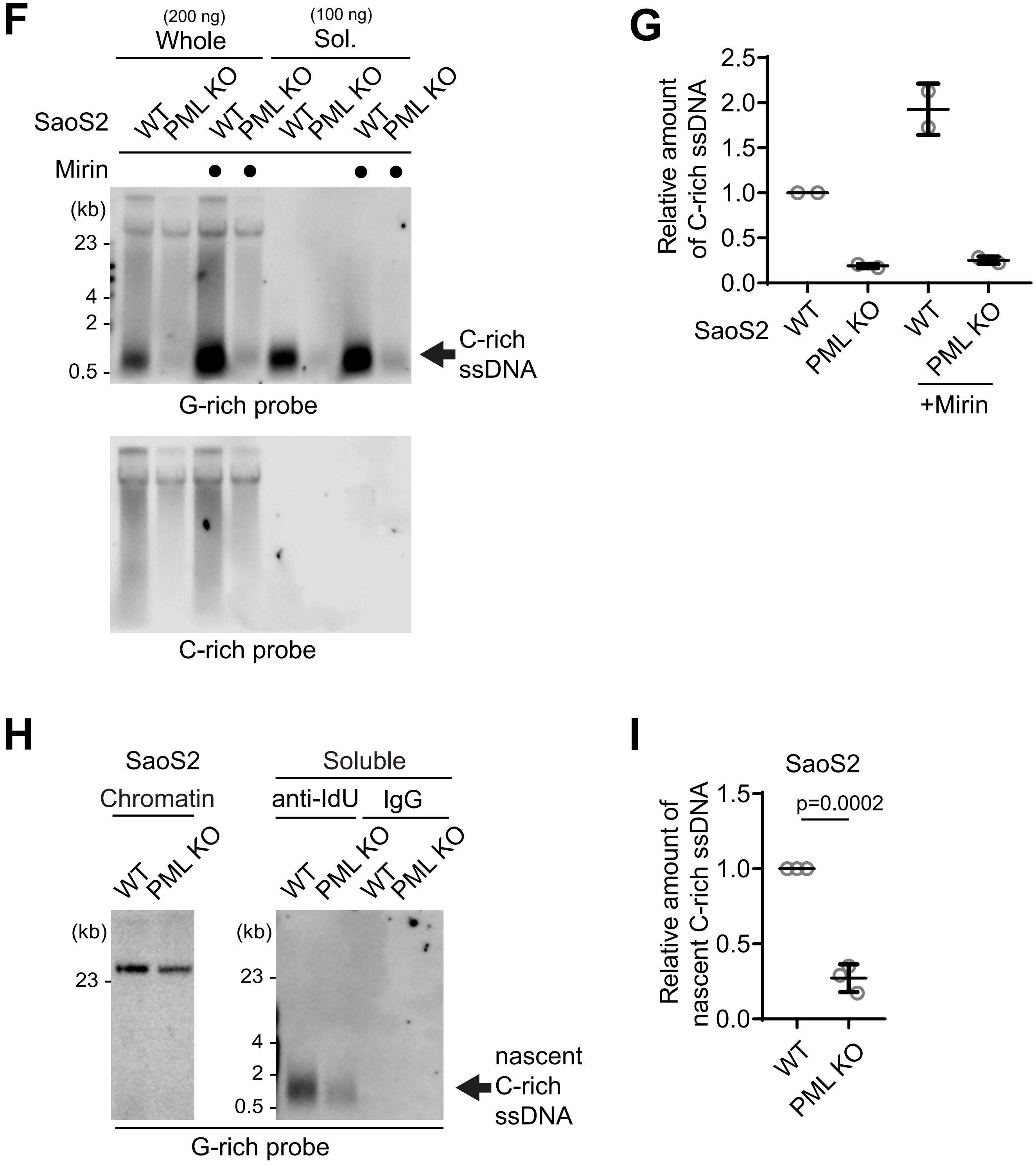
F
(200 ng)
Whole
(100 ng)
Sol.
SaoS2
WT
PML KO
WT
PML KO
WT
PML KO
WT
PML KO
Mirin
(kb)
23
4
2
0.5
C-rich
ssDNA
G-rich probe
C-rich probe
G
2.5
2.0
1.5
1.0
0.5
0.0
Relative amount
of C-rich ssDNA
SaoS2
WT
PML KO
WT
PML KO
+Mirin
H
SaoS2
Chromatin
Soluble
anti-IdU
IgG
WT
PML KO
WT
PML KO
WT
PML KO
(kb)
23
(kb)
23
4
2
0.5
nascent
C-rich
ssDNA
G-rich probe
I
SaoS2
1.5
1.0
0.5
0.0
Relative amount of
nascent C-rich ssDNA
p=0.0002
WT
PML KO

### Figure S4

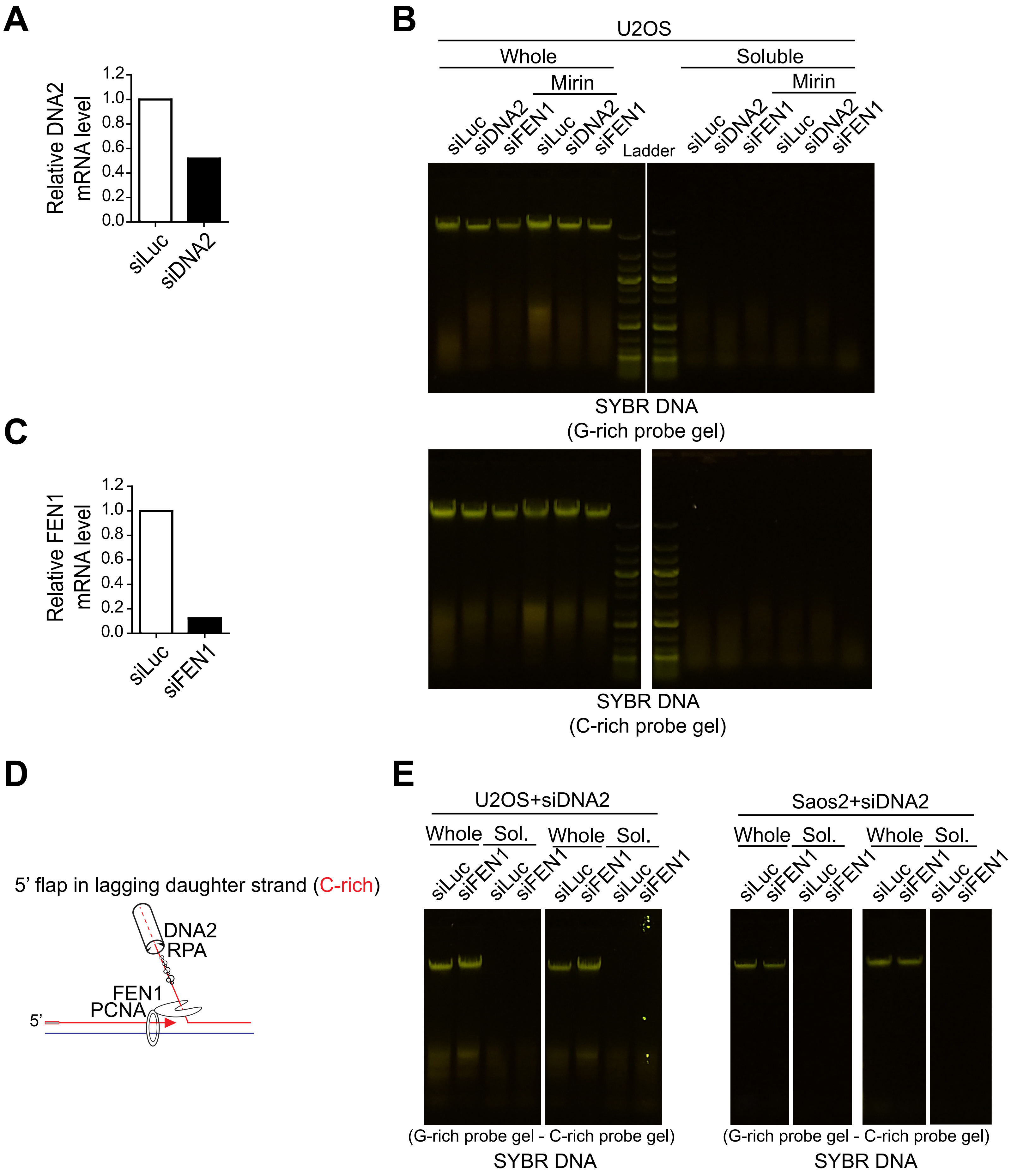
A
Relative DNA2 mRNA level
1.2
1.0
0.8
0.6
0.4
0.2
0.0
siLuc
siDNA2
B
U2OS
Whole
Soluble
Mirin
siLuc
siDNA2
siFEN1
Ladder
SYBR DNA
(G-rich probe gel)
SYBR DNA
(C-rich probe gel)
C
Relative FEN1 mRNA level
siFEN1
D
5' flap in lagging daughter strand (C-rich)
DNA2
RPA
FEN1
PCNA
5'
E
U2OS+siDNA2
Saos2+siDNA2
Whole
Sol.
(G-rich probe gel - C-rich probe gel)
SYBR DNA

### Figure S4

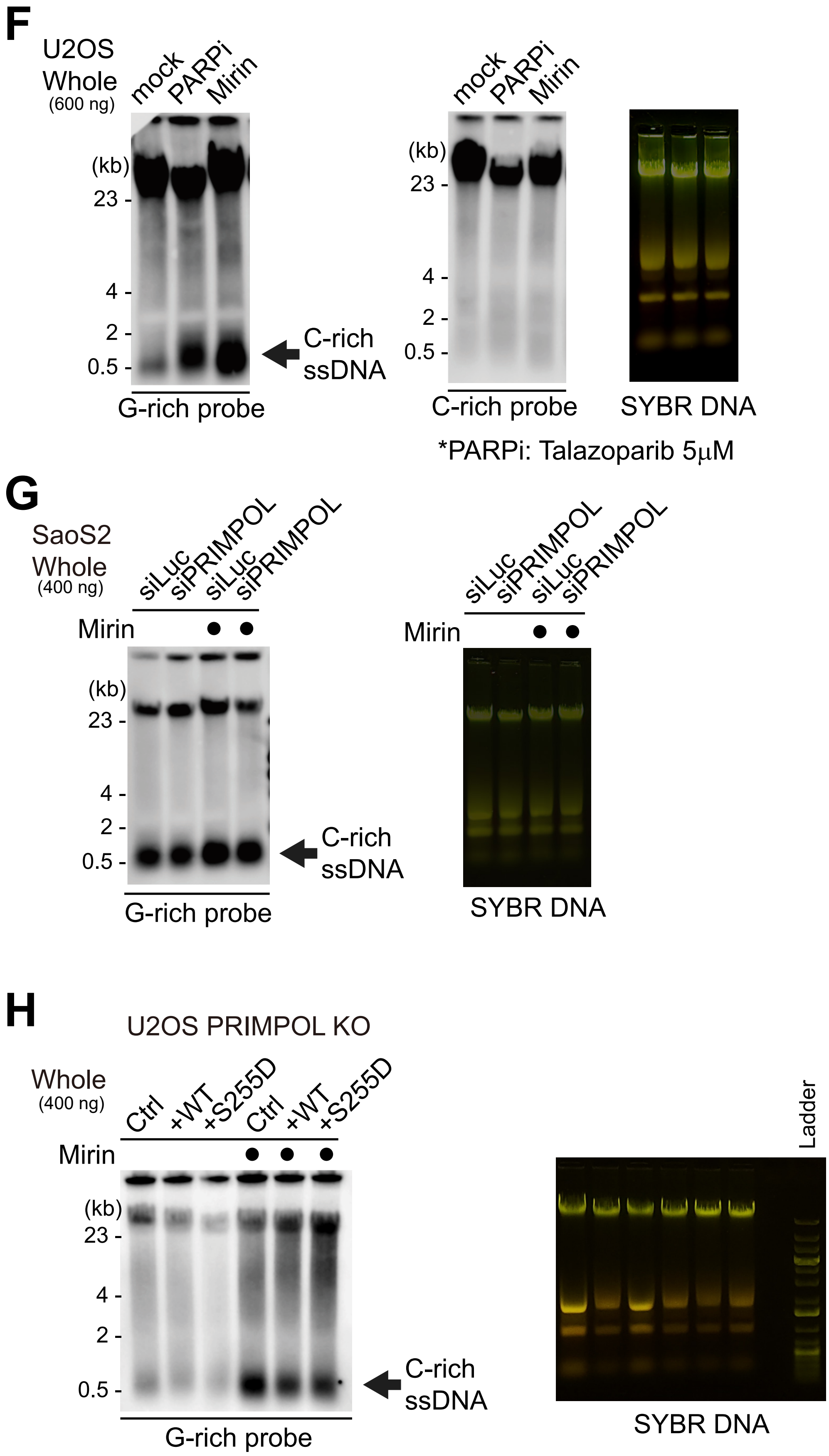
F
U2OS
Whole
(600 ng)
mock
PARPi
Mirin
(kb)
23 -
4 -
2 -
0.5 -
C-rich
ssDNA
G-rich probe
C-rich probe
SYBR DNA
*PARPi: Talazoparib 5μM
G
SaoS2
Whole
(400 ng)
siLuc
siPRIMPOL
siLuc
siPRIMPOL
Mirin
(kb)
23 -
4 -
2 -
0.5 -
C-rich
ssDNA
G-rich probe
SYBR DNA
H
U2OS PRIMPOL KO
Whole
(400 ng)
Ctrl
+WT
+S255D
Ctrl
+WT
+S255D
Mirin
(kb)
23 -
4 -
2 -
0.5 -
C-rich
ssDNA
G-rich probe
Ladder
SYBR DNA

Figure S5

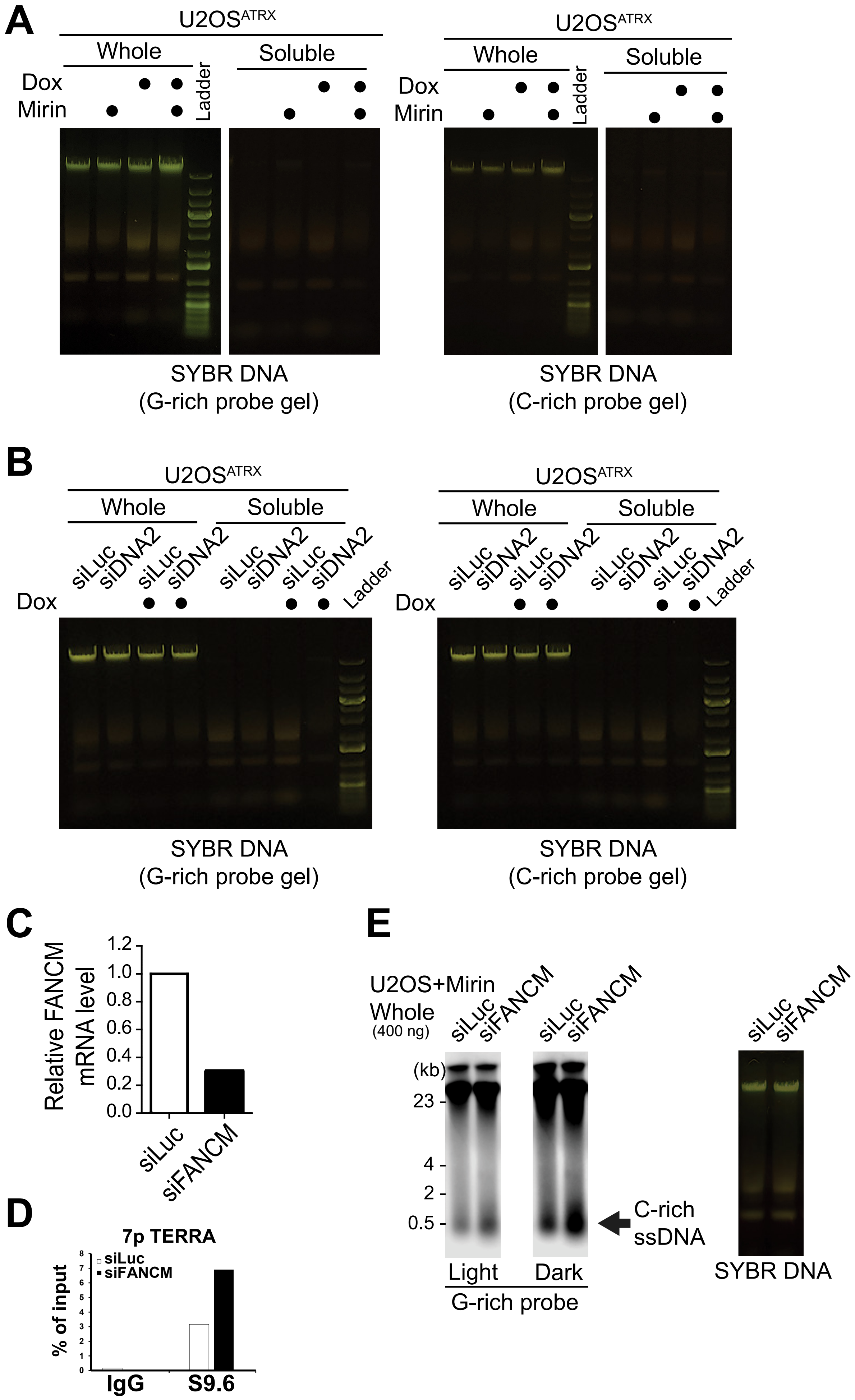

A
U2OS^ATRX
Whole
Soluble
Dox
Mirin
Ladder
SYBR DNA
(G-rich probe gel)
SYBR DNA
(C-rich probe gel)
B
Whole
Soluble
siLuc
siDNA2
Dox
Ladder
SYBR DNA
(G-rich probe gel)
SYBR DNA
(C-rich probe gel)
C
Relative FANCM mRNA level
1.2
1.0
0.8
0.6
0.4
0.2
0.0
siLuc
siFANCM
D
7p TERRA
siLuc
siFANCM
% of input
IgG
S9.6
E
U2OS+Mirin
Whole
(400 ng)
siLuc
siFANCM
(kb)
23
4
2
0.5
C-rich
ssDNA
Light
Dark
G-rich probe
SYBR DNA

### Figure S5

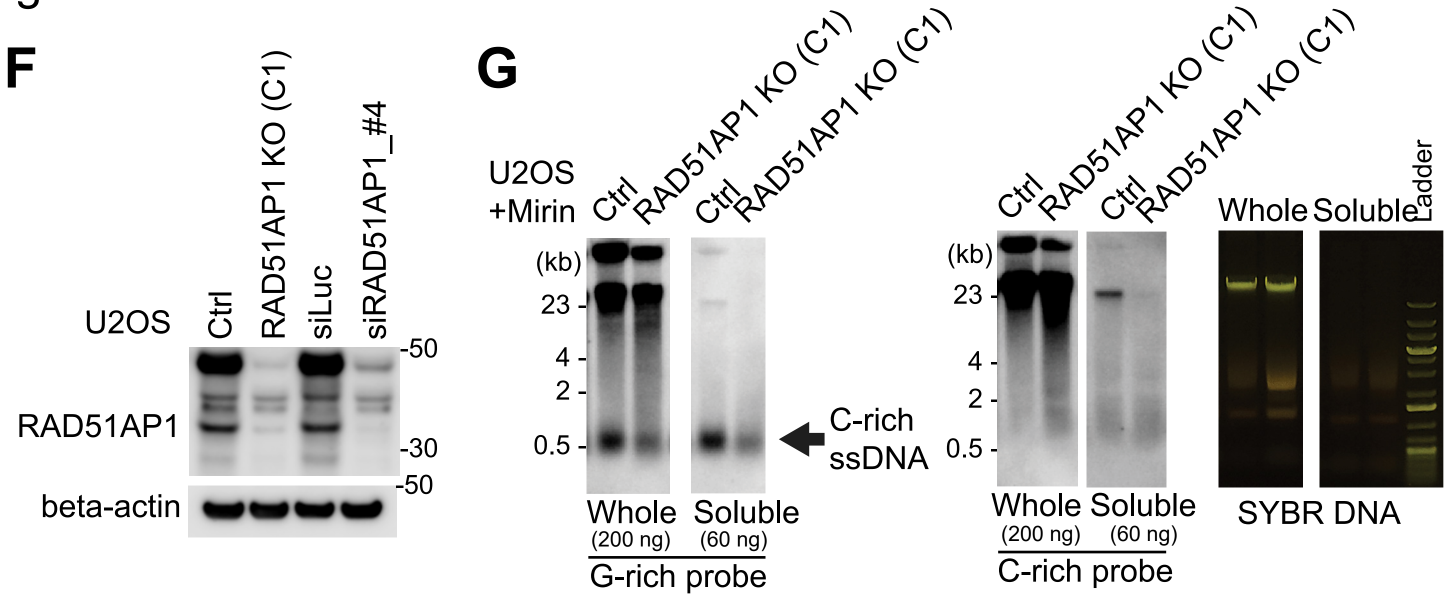

F
U2OS
Ctrl
RAD51AP1 KO (C1)
siLuc
siRAD51AP1_#4
RAD51AP1
beta-actin
-50
-30
-50
G
U2OS
+Mirin
Ctrl
RAD51AP1 KO (C1)
Ctrl
RAD51AP1 KO (C1)
(kb)
23
4
2
0.5
C-rich
ssDNA
Whole
(200 ng)
Soluble
(60 ng)
G-rich probe
C-rich probe
Whole
Soluble
Ladder
SYBR DNA


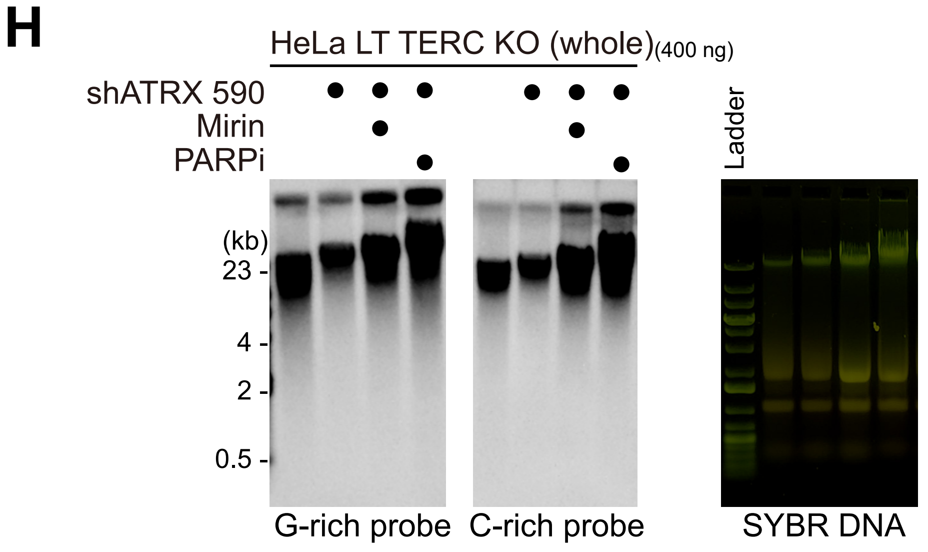

H
HeLa LT TERC KO (whole)(400 ng)
shATRX 590
Mirin
PARPi
(kb)
23
4
2
0.5
Ladder
G-rich probe
C-rich probe
SYBR DNA

### Figure S6

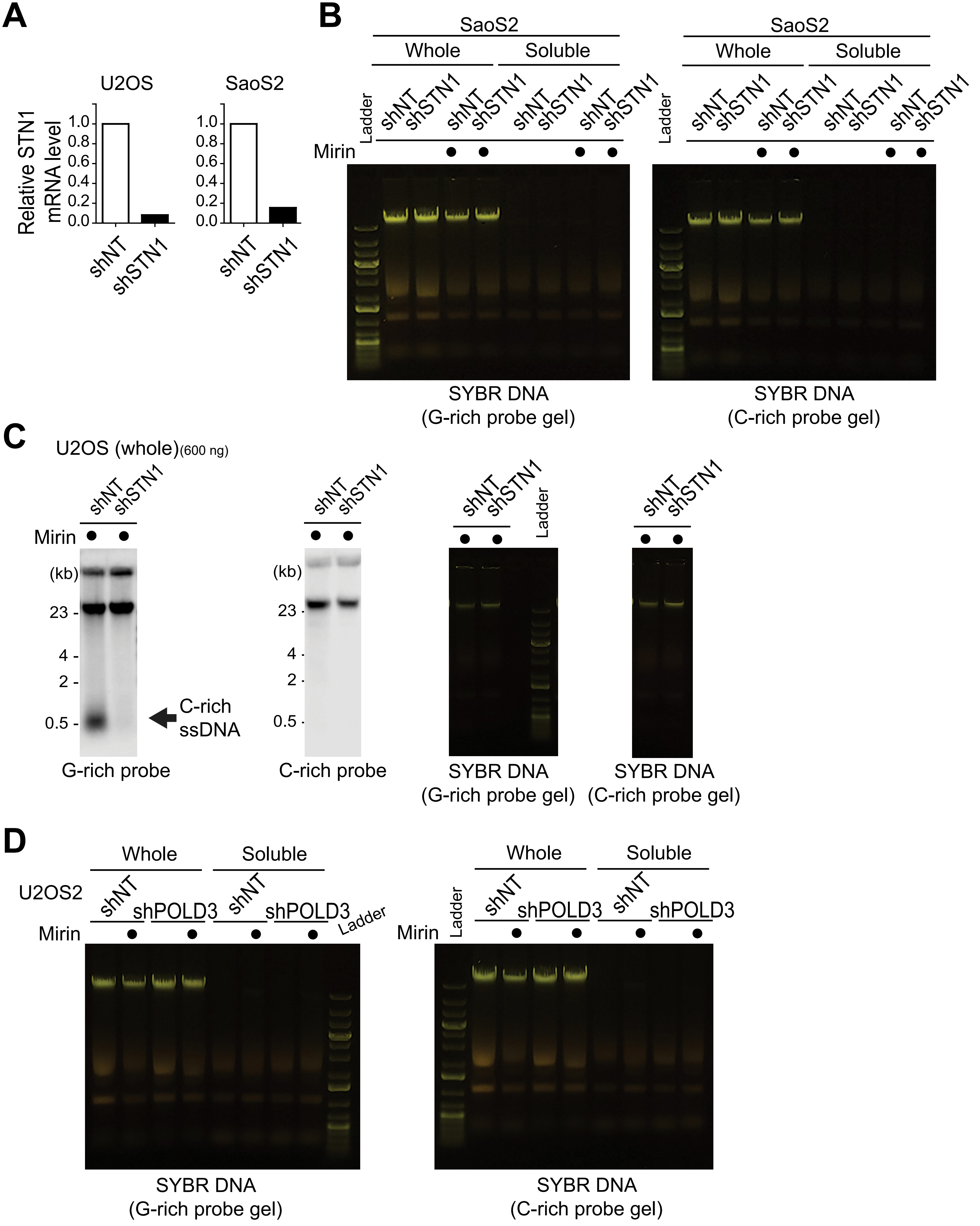

A
U2OS
SaoS2
Relative STN1 mRNA level
1.0
0.8
0.6
0.4
0.2
0.0
shNT
shSTN1
B
SaoS2
Whole
Soluble
Ladder
shNT
shSTN1
Mirin
SYBR DNA (G-rich probe gel)
SYBR DNA (C-rich probe gel)
C
U2OS (whole)(600 ng)
shNT
shSTN1
Mirin
(kb)
23
4
2
0.5
C-rich ssDNA
G-rich probe
C-rich probe
Ladder
SYBR DNA (G-rich probe gel)
SYBR DNA (C-rich probe gel)
D
Whole
Soluble
U2OS2
shNT
shPOLD3
Mirin
Ladder
SYBR DNA (G-rich probe gel)
SYBR DNA (C-rich probe gel)

Figure S6

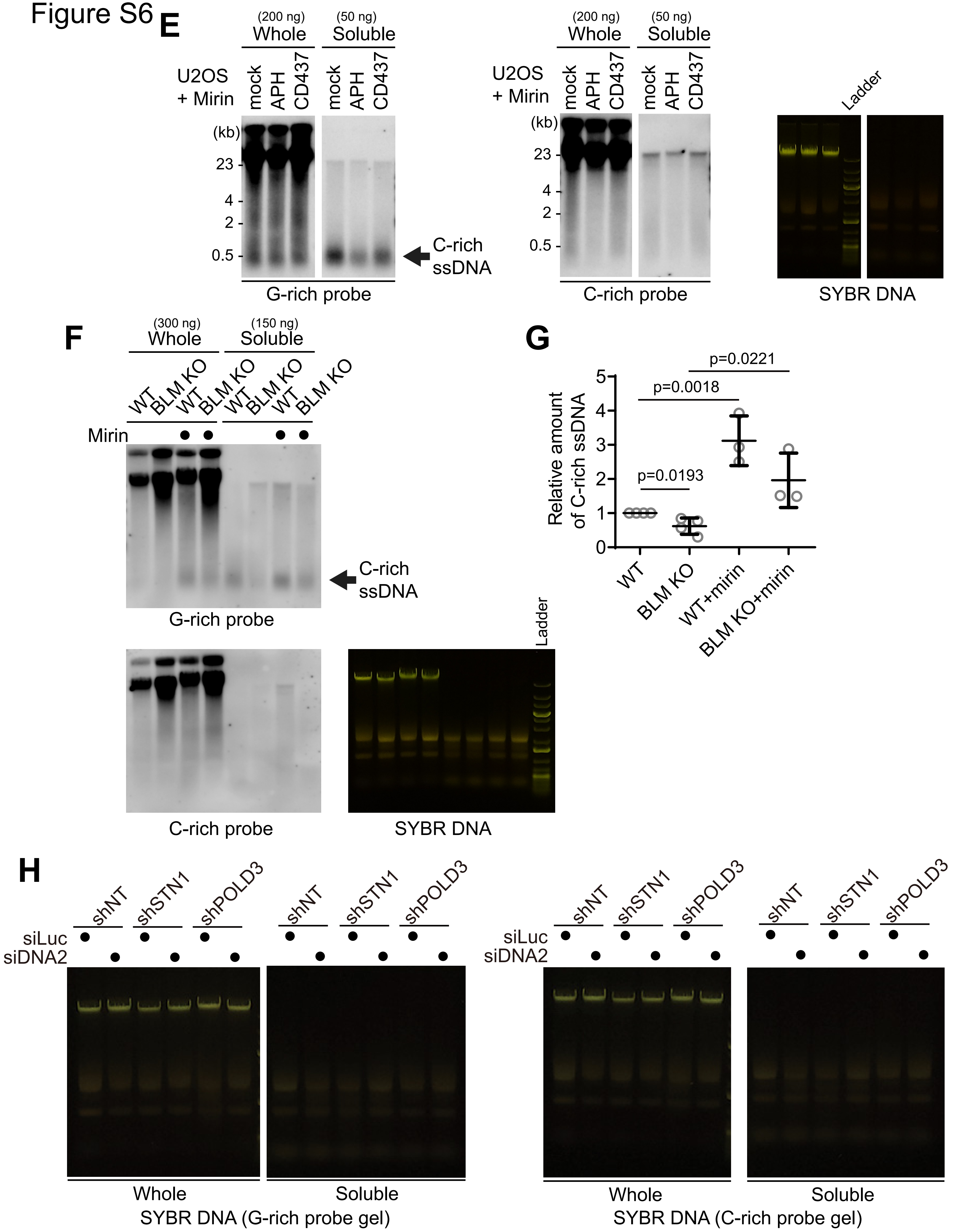
E
(200 ng)
Whole
(50 ng)
Soluble
U2OS
+ Mirin
mock
APH
CD437
mock
APH
CD437
(kb)
23
4
2
0.5
C-rich
ssDNA
G-rich probe
C-rich probe
Ladder
SYBR DNA
F
(300 ng)
Whole
(150 ng)
Soluble
WT
BLM KO
WT
BLM KO
WT
BLM KO
WT
BLM KO
Mirin
C-rich
ssDNA
G-rich probe
C-rich probe
SYBR DNA
G
Relative amount
of C-rich ssDNA
p=0.0221
p=0.0018
p=0.0193
5
4
3
2
1
0
WT
BLM KO
WT+mirin
BLM KO+mirin
H
shNT
shSTN1
shPOLD3
siLuc
siDNA2
Whole
Soluble
SYBR DNA (G-rich probe gel)
SYBR DNA (C-rich probe gel)
